# Comparative genomics reveals signatures of distinct metabolic strategies and gene loss associated with *Hydra* immortality

**DOI:** 10.64898/2026.02.11.705429

**Authors:** Kaoruko Nojiri, Koryu Kin, Akimasa Someya, Tetsuo Kon, Koto Kon-Nanjo, Hiroshi Shimizu, Kazuharu Arakawa, Etsuo A. Susaki

**Author notes:** Author for Correspondence: Etsuo A. Susaki, Department of Biochemistry and Systems Biomedicine, Juntendo University, Tokyo, Japan. These authors contributed equally to this work (Kaoruko Nojiri, Koryu Kin, Akimasa Someya).

## Abstract

*Hydra* is a freshwater cnidarian genus that provides a unique comparative model for aging research, contrasting the immortal *H. vulgaris* with the aging-inducible *H. oligactis.* Here, we report a high-quality, chromosome-level genome assembly of *H. vulgaris* strain AEP.JNIG. Our assembly is comparable in quality to existing resources, facilitating the analysis of genomic diversity across laboratory strains. Epigenomic profiling revealed that gene-body hypermethylation correlates with transcriptional stability and the suppression of spurious transcription in evolutionary conserved genes, suggesting an epigenetic mechanism for genomic integrity. Furthermore, comparative genomics demonstrated that while *Hydra* conserves fundamental *Hallmarks of Aging* pathways, the immortal *H. vulgaris* paradoxically lacks canonical anti-aging genes (e.g., Klotho, NAMPT) found in the aging-inducible *H. oligactis*. Instead, *H. vulgaris* exhibits a distinct metabolic signature related to mitochondrial energy production and NTP synthesis. Collectively, our comparative genomics results suggest multiple potential mechanisms associated with the *H. vulgaris* immortality and the aging traits of *H. oligactis*, providing novel targets for future functional studies.

**Significance statement:** Why do some organisms age while others appear not to? The freshwater animal *Hydra* provides a unique opportunity to investigate this question, as closely related species display contrasting aging phenotypes. We generated a high-quality genome assembly for a new strain of a non-aging species and conducted comparative analyses with related strains and an aging species. Even closely related strains can accumulate substantial genetic divergence over time, and stable DNA modification patterns were associated with consistent gene activity, suggesting a mechanism that may help maintain cellular balance. Surprisingly, several well-known longevity genes are present in the aging species but absent in the non-aging one. This suggests that extended lifespan may not simply depend on possessing more “anti-aging” genes, but instead may reflect differences in how core biological processes are organized. Our study provides new insights into the genetic basis of aging and highlights *Hydra* as a powerful model for understanding longevity.

## Introduction

*Hydra* is a freshwater cnidarian that can be maintained in the laboratory in mass culture (Klimovich et al. 2019; Galliot 2012; Loomis & Lenhoff 1956). Based on its distinctive traits including potent regenerative capacity and both sexual and asexual modes of reproduction, *Hydra* has been widely studied in fields such as stem cell biology, regeneration, and aging (Holstein 2023; Vogg et al. 2019; Tomczyk et al. 2015). In particular, *Hydra vulgaris* is one of the few organisms in which negligible senescence has been demonstrated by long-term population monitoring of mortality and fertility (Martínez 1998; Schaible et al. 2015). Aging organisms exhibit progressive decline in physiological function and accumulation of pathological changes over time, characterized by an increase in mortality rate (Gompertz curve) (Finch et al. 1990; Jones et al. 2014). On the other hand, *H. vulgaris* maintains a flat Gompertz curve (Schaible et al. 2015), suggesting its immortality. This remarkable trait is thought to be supported by the continuous self-renewal capacity of its stem cell populations, regulated by genes including FoxO and TCF/β-catenin (Boehm et al. 2012; Hemmrich et al. 2012; Gufler et al. 2018). Interestingly, *H. oligactis*, another species within the *Hydra* genus, shows increased mortality and physiological deterioration after sexual reproduction when transferred to 10 °C (Yoshida et al. 2006). This inducible aging phenotype has been attributed to the depletion of interstitial stem cells (ISCs) (Sun et al. 2020; Tomczyk et al. 2020).

*Hydra* also constitutes an apparent exception to evolutionary theories of aging. These theories generally propose that organismal aging reflects an evolutionary trade-off that favors an energy-saving strategy, in which ensuring germline accuracy is essential, whereas maintaining somatic cell integrity is often a dispensable expenditure (Kirkwood 1977; Hamilton 1966). Therefore, whereas many long-lived species delay reproduction until later developmental stages, a pattern consistent with such theoretical predictions, *Hydra* attains sexual maturity early in life yet retains an extended lifespan (Schaible et al. 2015).

What might be the origin of these unique aging traits in *Hydra*? While organismal aging is generally considered a complex process involving both genetic and environmental factors, empirical evidence suggests that genetic components represent the primary determinant. Multiple findings support this proposition. First, maximum lifespan and aging patterns described by Gompertz curves show species-specific differences (Jones et al. 2014). Second, multiple aging-related genes and pathways have been identified, such as Klotho, progeroid syndrome-related genes including DNA repair and nuclear integrity, and NAD^+^-related pathways (Nabeshima 2002; Lombard et al. 2005). The *Hallmarks of aging* provides a comprehensive framework that summarizes genes and biological processes in aging (López-Otín et al. 2023). Third, tissue and organismal aging is closely associated with the resilience of tissue stem cells, which is also determined by genomic information (Rando & Jones 2021). Lastly, a recent study demonstrated that the heritability of intrinsic life span is approximately 50%, suggesting that genetic control over longevity is comparable to other complex traits (Shenhar et al. 2026). Therefore, comparative genomic analysis may provide critical insights into the genetic elements underlying the immortality observed in *H. vulgaris*.

While high-quality whole-genome assembly for the *H. vulgaris* strain AEP (North American strain) (Cazet et al. 2023), originally established in North America (Martínez et al. 2010), provides useful references for comparative genomics, they do not capture the full extent of intra-species genomic variation. Such variation may arise from spontaneously occurring mutants in laboratory cultures. Indeed, independent maintenance of laboratory populations or colonies can generally lead to genomic divergence across species (Chebib et al. 2021; Farslow et al. 2015). In *H. vulgaris* 105 (formerly *H. magnipapillata*) maintained by asexual budding, the stem cells have accumulated mutations at even slightly higher rates than those in mammals, with some under positive selection (Sahm et al. 2024). Distinct types of transposable elements, such as the CR1 LINE element, are inserted into stem cell genomes, further contributing to genomic divergence (Kon-Nanjo et al. 2025; Wong et al. 2019). These suggest that the *Hydra* genome can change over long-term cultures in laboratories, even when the animals reproduce asexually.

To avoid risks of overlooking population-specific variants that can affect phenotypic traits, at least two high-quality reference genomes from independently maintained *H. vulgaris* populations are desirable. To address this, we sequenced and assembled a chromosome-level genome for the *H. vulgaris* AEP strain maintained at the National Institute of Genetics, Mishima, Japan (hereafter *H. vulgaris* strain AEP.JNIG) (Kawaida et al. 2010), thereby expanding available genomic resources for this species. We verified that *H. vulgaris* AEP.JNIG forms a monophyletic group with other *H. vulgaris* strains, including AEP and 105, and is the most closely related to AEP, while chromosomal analyses revealed substantial structural differences, confirming genome-level divergence among populations. Building on these data, we conducted a comparative analysis of aging-related genes and pathways using the genomes of three *H. vulgaris* strains and the aging species *H. oligactis* (Cazet et al. 2023). By comparing signaling pathways and genes tied to the *Hallmarks of Aging* (López-Otín et al. 2023), we identified gene sets that differ between aging and non-aging species. From these comparisons, we characterized genomic differences between *H. vulgaris* and representative aging animals, including *H. oligactis*, and identified candidate genes potentially associated with *H. vulgaris* immortality.

## Results

### Chromosome-level genome assembly of *H. vulgaris* AEP.JNIG

The *H. vulgaris* population maintained at the National Institute of Genetics, Mishima, Japan (*H. vulgaris* AEP.JNIG, Fig 1A) was investigated. We independently sequenced and assembled the *H. vulgaris* AEP.JNIG genome as part of the Japanese *Hydra* resource; during the course of our work, a genome assembly of an *H. vulgaris* AEP strain maintained in the US was published (Cazet et al. 2023). The availability of both assemblies allowed us to evaluate the quality of the AEP.JNIG genome and to perform population-level comparisons of genomic variation.

**Figure 1.**
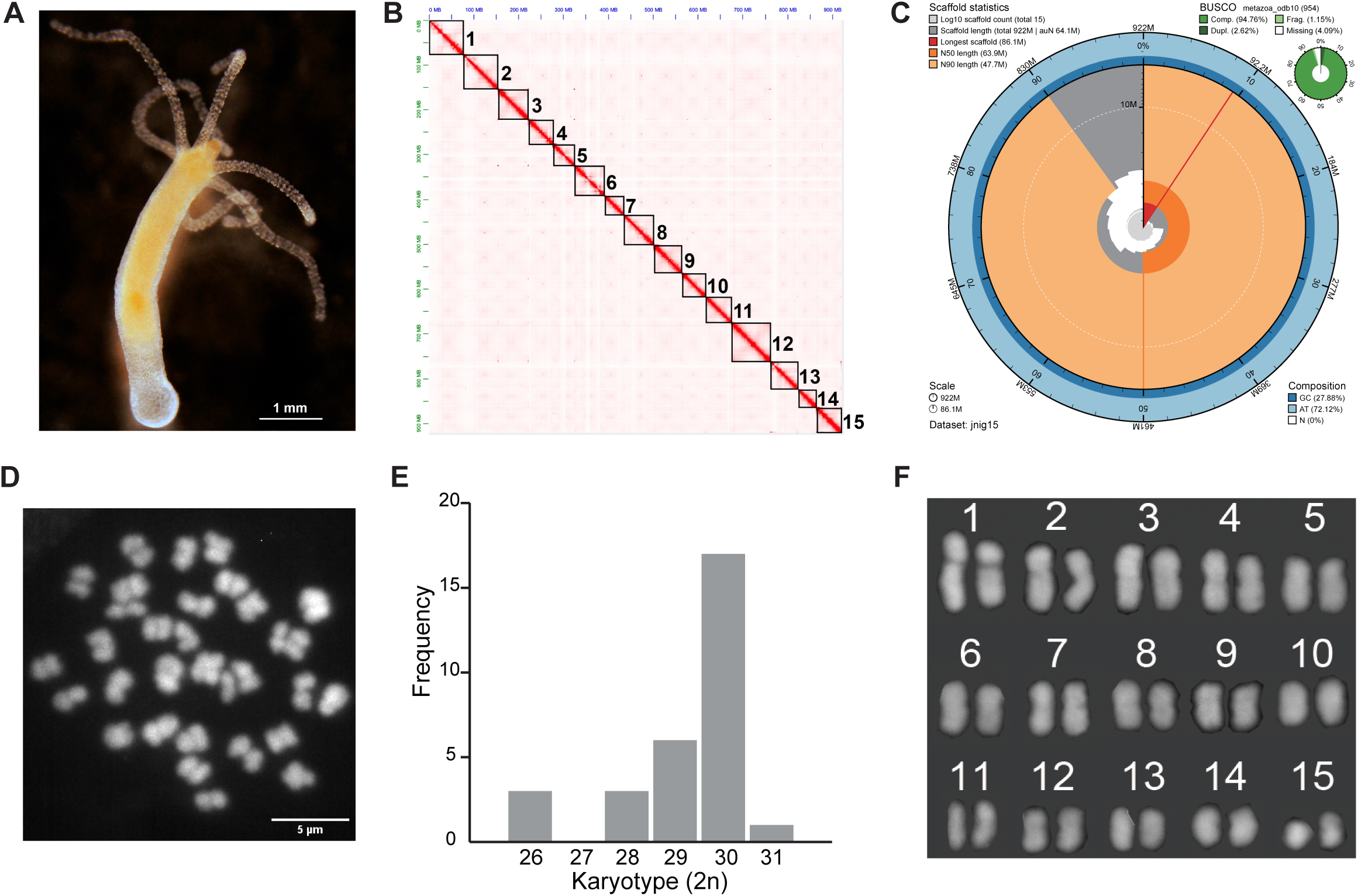
Chromosome-level genome assembly of *H. vulgaris* AEP.JNIG. **(A)** Picture of cultured *H. vulgaris* AEP.JNIG. Scale bar: 1 mm. **(B)** Hi-C contact map, showing the 15 chromosome-level scaffolds of *H. vulgaris* AEP.JNIG. Darker red indicates genomic regions with higher contact frequencies, reflecting closer spatial proximity. **(C)** Snail plot demonstrating the quality of our genome assembly. **(D)** A representative image of metaphase chromosomes in *H. vulgaris* AEP.JNIG stained with DAPI and observed with a scanning disk confocal microscope. Scale bar: 5 μm. **(E)** Histogram of karyotypes observed in this study. Chromosome counts show a distinct peak at 2n = 30. The absence of nuclei with 2n > 32 supports a chromosome number of 30, consistent with our and other genome assembly data. **(F)** Karyogram of *Hydra vulgaris* AEP.JNIG showing 15 chromosomes. **Alt text:** Composite figure with panels A–F illustrating the chromosome-level genome assembly and karyotype of *Hydra vulgaris* AEP.JNIG. A is a photograph of a cultured *H. vulgaris* polyp. B is a Hi-C contact map displayed in red and white colors showing 15 chromosome-scale scaffolds as strong diagonal blocks indicating high contact frequencies within chromosomes. C is a circular “snail plot” summarizing genome assembly statistics, including BUSCO completeness of 94.7%. D is a fluorescence microscopy image of metaphase chromosomes stained with DAPI, showing a diploid set of chromosomes. E is a histogram of chromosome counts with a clear peak at 2n = 30. F is a karyogram showing 15 chromosome pairs arranged from longest to shortest.

The whole genome sequence of *H. vulgaris* AEP.JNIG genome was initially collected as the assemblies of Illumina paired-end reads and Nanopore long reads. We obtained 82,804,328,738 bp of paired-end short reads and 168,380,136,564 bp of Nanopore long reads with an N50 read length of 3,493 bp (Table 1). *De novo* assembly was performed using these sequences with Phased Error Correction and Assembly Tool (PECAT) (Nie et al. 2024) (see Materials and Methods for details). The initial genome assembly had a total size of 954,340,183 bp, consisting of 248 contigs. The longest contig was 61,504,050 bp, the contig N50 was 9,577,232 bp, the contig L50 was 30, and the GC content was 27.94%. (Table 2).

**Table 1.** Summary of the genome sequencing.

|  | Total read | Total base | Read length (N50) |
| --- | --- | --- | --- |
| WGS (Illumina) | 548,373,038 | 82,804,238,738 | 2x150 |
| WGS (Nanopore) | 63,239,530 | 168,380,136,564 | 3493.0 |
| Hi-C (Illumina) | 796,383,332 | 119,457,499,800 | 2x150 |

To construct a chromosome-level genome assembly, 398,191,666 Hi-C read pairs were generated from the same clone. We scaffolded the initial genome assembly using the Hi-C reads with HiC-pro (Servant et al. 2015) and YaHS (Zhou et al. 2022) (see Materials and Methods for details). Fifteen chromosome-level scaffolds were obtained, each showing strong intra-chromosomal contact signals in the Hi-C contact map (Fig 1B). The total size of the final assembly was 922,069,947 bp, the largest scaffold was 86,117,336 bp, scaffold N50 was 63,895,935 bp, the contig L50 was 7, and the GC content was 27.88% (Table 2).

**Table 2.** Assembly statistics of the H. vulgaris genome.

| QUAST v5.3.0 | Initial assembly AEP.JNIG | Final Assembly AEP.JNIG | Published Assembly AEP (Cazet *et al.* 2023) |
| --- | --- | --- | --- |
| Total length | 954,340,183 | 922,069,947 | 900,935,055 |
| Scaffolds | 248 | 15 | 15 |
| Longest scaffold | 61,504,050 | 86,117,336 | 85,887,368 |
| N50 | 9,577,232 | 63,895,935 | 59,555,380 |
| L50 | 30 | 7 | 7 |
| GC content | 27.94 | 27.88 | 27.77 |

We evaluated the accuracy and completeness of the final genome assembly by searching for sets of BUSCO (v 5.8.3) (Manni et al. 2021) using the metazoa odb10 database (Kriventseva et al. 2019). We identified 904 complete BUSCO genes out of 954 (94.7%), consisting of 879 single-copy genes (92.1%) and 25 duplicated genes (2.6%). 11 BUSCO genes were identified as fragmented (1.2%), and 39 BUSCO genes were missing (4.1%) (Fig 1C and Table 3). These assembly results are comparable to those for the *H. vulgaris* AEP genome (Cazet et al. 2023) (Tables 2 and 3), supporting the sufficient quality of our genome assembly dataset. We further created gene models based on this chromosome-level assembly and the transcriptome using BREAKER2 (Brůna et al. 2021) and evaluated BUSCO scores. This analysis identified 891 complete BUSCO genes out of 954 (93.4%), consisting of 772 single-copy genes (80.9%) and 119 duplicated genes (12.5%). 14 BUSCO genes were fragmented (1.5%), and 49 BUSCO genes were missing (5.1%). Again, these results are also comparable to those for the AEP genome (Cazet et al. 2023) (Table 4).

**Table 3.**
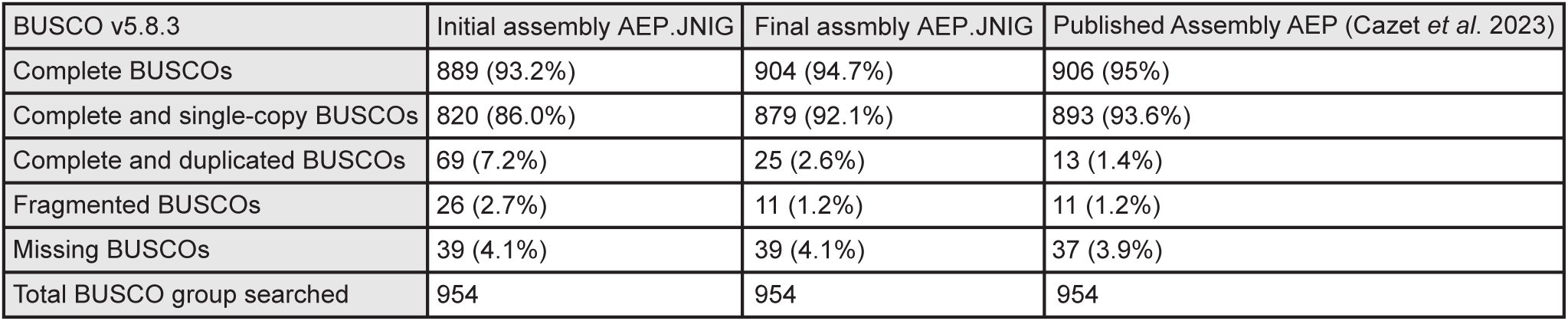
BUSCO score from the genome assembly.

| BUSCO v5.8.3 | Initial assembly AEP.JNIG | Final assmblly AEP.JNIG | Published Assembly AEP (Cazet *et al.* 2023) |
| --- | --- | --- | --- |
| Complete BUSCOs | 889 (93.2%) | 904 (94.7%) | 906 (95%) |
| Complete and single-copy BUSCOs | 820 (86.0%) | 879 (92.1%) | 893 (93.6%) |
| Complete and duplicated BUSCOs | 69 (7.2%) | 25 (2.6%) | 13 (1.4%) |
| Fragmented BUSCOs | 26 (2.7%) | 11 (1.2%) | 11 (1.2%) |
| Missing BUSCOs | 39 (4.1%) | 39 (4.1%) | 37 (3.9%) |
| Total BUSCO group searched | 954 | 954 | 954 |

**Table 4.** BUSCO score from the gene model.

| BUSCO v5.8.3 | Final assmblly AEP.JNIG | Published Assembly AEP (Cazet *et al.* 2023) |
| --- | --- | --- |
| Complete BUSCOs | 891 (93.4%) | 905 (94.9%) |
| Complete and single-copy BUSCOs | 772 (80.9%) | 705 (73.9%) |
| Complete and duplicated BUSCOs | 119 (12.5%) | 200 (21.0%) |
| Fragmented BUSCOs | 14 (1.5%) | 14 (1.5%) |
| Missing BUSCOs | 49 (5.1%) | 35 (3.7%) |
| Total BUSCO group searched | 954 | 954 |

To further confirm the chromosome-level integrity of our AEP strain, we also performed karyotyping (Fig 1D-F). Our microscopic observations predominantly indicate the karyotype of 2n = 30, with a few exceptions likely due to artifacts during sample preparations (Fig 1E, S1 Fig). Importantly, the absence of any karyotypes with 2n > 32 strongly suggests that *H. vulgaris* AEP.JNIG does not possess extra chromosome pairs compared to other *H. vulgaris* strains. Therefore, we conclude that the *H. vulgaris* AEP.JNIG has a karyotype of 2n = 30 (Fig 1F).

### Comparison of *H. vulgaris* AEP.JNIG genome with AEP genome revealed overall conservation of chromosomal structure

To determine the phylogenetic relationship between our strain and the other *Hydra* strains, we performed species tree reconstruction using a large-scale phylogenomic dataset (a multiple sequence alignment of 981,632 amino acid positions from 954 proteins) from 31 metazoan organisms including those of two other *H. vulgaris* strains and 2 holozoan protists (Cazet et al. 2023; Kon-Nanjo et al. 2025; Chapman et al. 2010) (see Materials and Methods for details) (Fig 2A). The resulting species tree robustly placed *H. vugaris* AEP.JNIG as the closest relative of the *H. vulgaris* AEP strain. Furthermore, in order to see whether there is any strain closer to AEP.JNIG than AEP, we extracted four genes (*COI*, *COII*, *EF1α*, *Cnnos1*) from the AEP.JNIG genome and made a phylogenetic tree with other strains using a dataset previously published (Kawaida et al. 2010). In all cases, *H. vulgaris* AEP.JNIG formed a monophyletic group with *H. vulgaris* AEP (S2 Fig), further corroborating a close phylogenetic relationship between *H. vulgaris* AEP.JNIG and *H. vulgaris* AEP.

**Figure 2.**
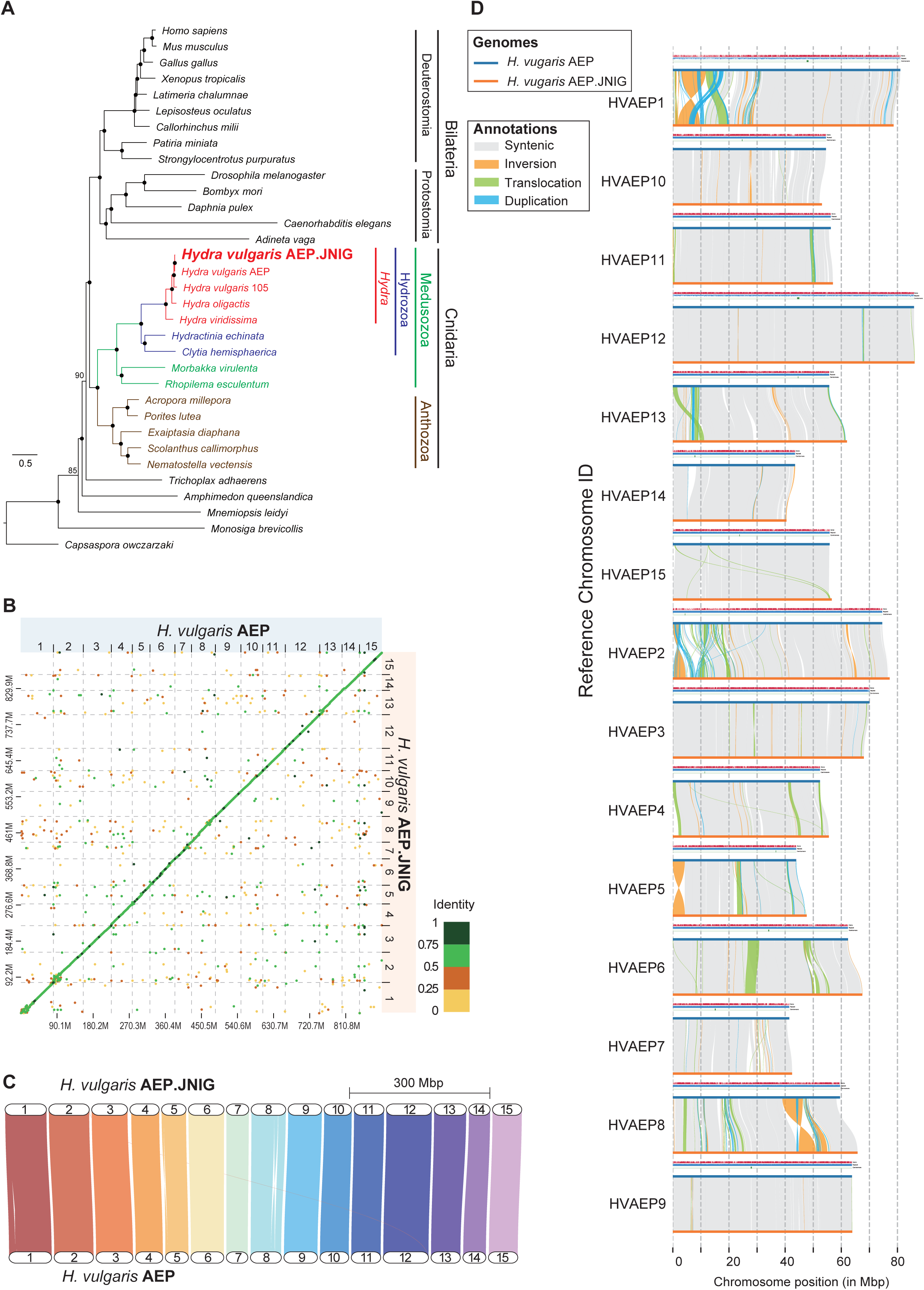
Comparison of *H. vulgaris* AEP and *H. vulgaris* AEP.JNIG genome assemblies. **(A)** Maximum likelihood phylogenetic analysis of a phylogenomic dataset constructed from 33 holozoan proteomes shows that *H. vulgaris* AEP and *H. vulgaris* AEP.JNIG form a monophyletic clade. Nodes marked with black dots (●) indicate 100% ultrafast bootstrap support, while other nodes were labelled with their corresponding ultrafast bootstrap support values. The scale bar represents 0.5 amino acid substitutions per site. **(B)** A dot-plot created with D-GENIES shows that the genome of *H. vulgaris* JNIG (on the y-axis) aligns well with that of *H. vulgaris* AEP genome (Cazet et al. 2023) (on the x-axis, indicating a high genomic similarity between the two strains. The color indicates the identity score as shown in the legend on the right. **(C)** Gene synteny plot generated by GENESPACE showing conserved synteny of orthologous genes between the AEP and AEP.JNIG genomes. The AEP.JNIG chromosomes are shown at the top and the AEP chromosomes at the bottom. Chromosome numbering and sequence strand orientation were standardized to match previously published genome assemblies. **(D)** Visualization of syntenic regions and structural rearrangements between chromosomes from *H. vulgaris* AEP and *H. vulgaris* AEP.JNIG genomes. Annotation tracks for four genomic features (number of SNPs, genes, repetitive regions and centromeres) were included using optional parameters above the lines showing the chromosomes of each strain. In the annotation tracks, higher values indicate greater density. **Alt text**: Composite figure with panels A–D comparing the genome assemblies of *Hydra vulgaris* AEP and AEP.JNIG. A is a phylogenetic tree based on a dataset of 33 holozoan proteomes showing that AEP and AEP.JNIG form a monophyletic clade, with bootstrap support values indicated at the nodes. B is a genome alignment dot plot with the AEP genome on the x-axis and the AEP.JNIG genome on the y-axis, showing strong diagonal alignments indicating high genomic similarity. C is a gene synteny plot showing conserved syntenic relationships between chromosomes of the AEP.JNIG genome and the AEP genome. D is a visualization of syntenic regions and structural rearrangements between chromosomes of the two genomes, with annotation tracks showing the density of SNPs, genes, repetitive regions, and centromeres.

To characterize the degree of genetic divergence between AEP and AEP.JNIG, we compared their genomes using D-GENIES (Cabanettes & Klopp 2018) and GENESPACE (Lovell et al. 2022) (Fig 2B and 2C). The dot plot from the genome alignment between these two genome sequences by D-GENIES revealed prominent diagonal lines, indicating a high degree of collinearity between them (Fig 2B). The ribbon plot from the gene syntenic analysis result by the GENESPACE demonstrated overall correspondence at the chromosomal level (Fig 2C). We then used plotsr (Goel & Schneeberger 2022) and SyRI (Goel et al. 2019) to visualize the detailed genomic structural variations such as inversions, translocations, and duplications, and to quantify both structural and sequence-level differences including SNPs and indels. In total, 954 syntenic regions were identified, covering 796 Mb of the AEP assembly and 789 Mb of the AEP.JNIG assembly. In addition, 360 inversions (31.5 Mb) and 660 translocations (31.5 Mb) were also detected, along with numerous duplication events (239 in the AEP and 2,073 in the AEP.JNIG) (Fig 2D and S3A Fig). At the sequence level, 1,833,472 SNPs, 648,211 insertions (5.4 Mb in total), and 261,207 deletions (4.7 Mb in total) were identified (S3A Fig). Together, these results indicate that overall chromosomal organization is largely conserved between the two strains, despite substantial accumulation of sequence-level variation. Although we observed structural variations between the AEP and AEP.JNIG assemblies, we cannot entirely rule out the possibility that some of these differences arise from misassemblies in either genome. However, given the strong signals observed in the Hi-C contact maps for both assemblies, such artifacts are likely minimal.

We also examined the transposable element (TE) landscape of the *H. vulgaris* AEP.JNIG assembly using Earl Grey (v6.3.3) (Baril et al. 2024). Repetitive elements comprised 78.4% of the total genome (Table 5). DNA transposons were the most abundant class, representing 35.8% of the genome, and LINEs accounted for 15.2%. By contrast, LTR and Rolling-circle elements were relatively rare (1.0% and 1.0%, respectively). For direct comparison, we applied the same analysis to the *H. vulgaris* AEP assembly. The overall repeat content (77.7%) and element composition were similar between the two populations, although LINEs and Peneopes were slightly more abundant in AEP.JNIG (Table 6), indicating general consistency between the two assemblies. Despite slight differences in subfamily composition, TE landscape comparisons between the AEP.JNIG and AEP assemblies showed overall consistency (S3B and S3C Figs).

**Table 5.** Composition of TEs identified by Earl Grey in the genomes of H. vulgaris AEP.JNIG.

| TE Classification | Coverage (bp) | Copy number | % Genome coverage | Genome size | TE family count |
| --- | --- | --- | --- | --- | --- |
| DNA | 330,098,105 | 597,308 | 35.8 | 922,069,947 | 597,307 |
| Rolling Circle | 8,793,916 | 19,836 | 0.954 | 922,069,947 | 19,836 |
| Penelope | 23,760,322 | 68,248 | 2.58 | 922,069,947 | 68,248 |
| LINE | 140,003,727 | 267,317 | 15.2 | 922,069,947 | 267,317 |
| LTR | 9,088,772 | 10,622 | 0.986 | 922,069,947 | 10,622 |
| Other (Simple Repeat, Microsatellite, RNA) | 54,617,458 | 421,636 | 5.92 | 922,069,947 | 421,636 |
| Unclassified | 156,717,139 | 504,751 | 17.0 | 922,069,947 | 504,750 |
| **Repeat** | 723,079,439 | NA | 78.4 | 922,069,947 | NA |
| **Non-Repeat** | 198,990,508 | NA | 21.6 | 922,069,947 | NA |

**Table 6.**
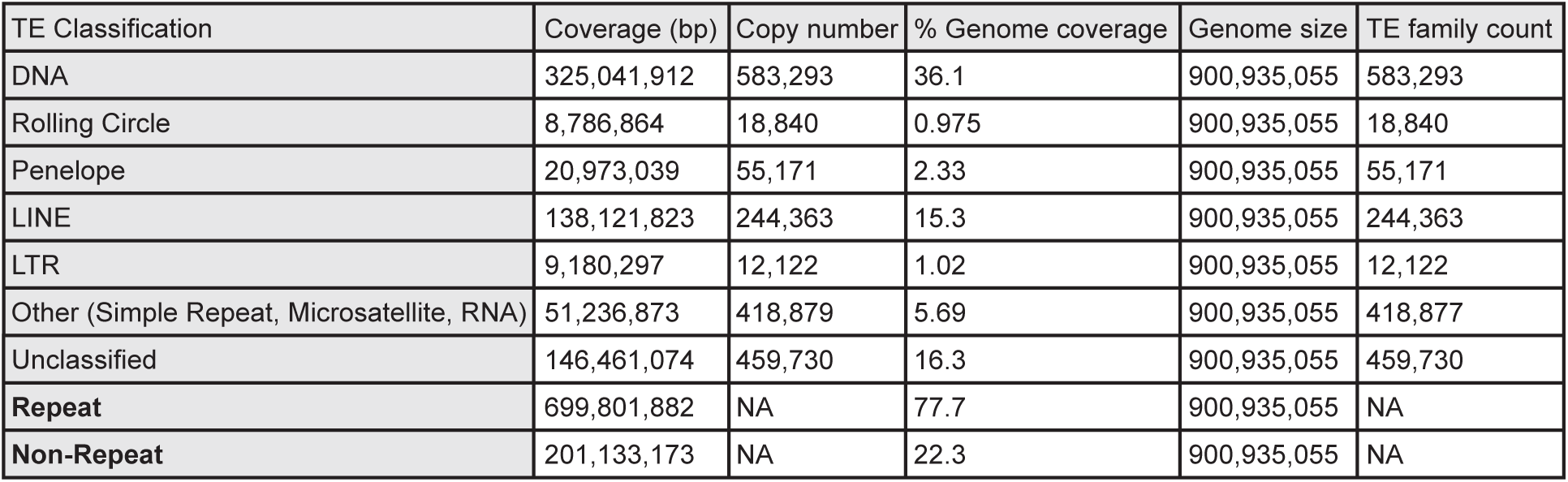
Composition of TEs identified by Earl Grey in the genomes of H. vulgaris AEP.

### DNA Methylation within gene bodies contributes to transcriptional stability (homeostasis) of conserved genes in *H. vulgaris*

Leveraging the methylation signal in the Nanopore reads, we examined genome-wide mCpG distribution in *H. vulgaris* AEP.JNIG using Dorado (v0.7.1) and Modkit (v0.4.3) (see Materials and Methods). A mosaic methylation pattern was observed across the genome, with stably methylated and unmethylated regions interspersed. Centromeric regions identified by quarTeT (v1.2.5) (Lin et al. 2023) had low DNA methylation (Fig 3A).

**Figure 3.**
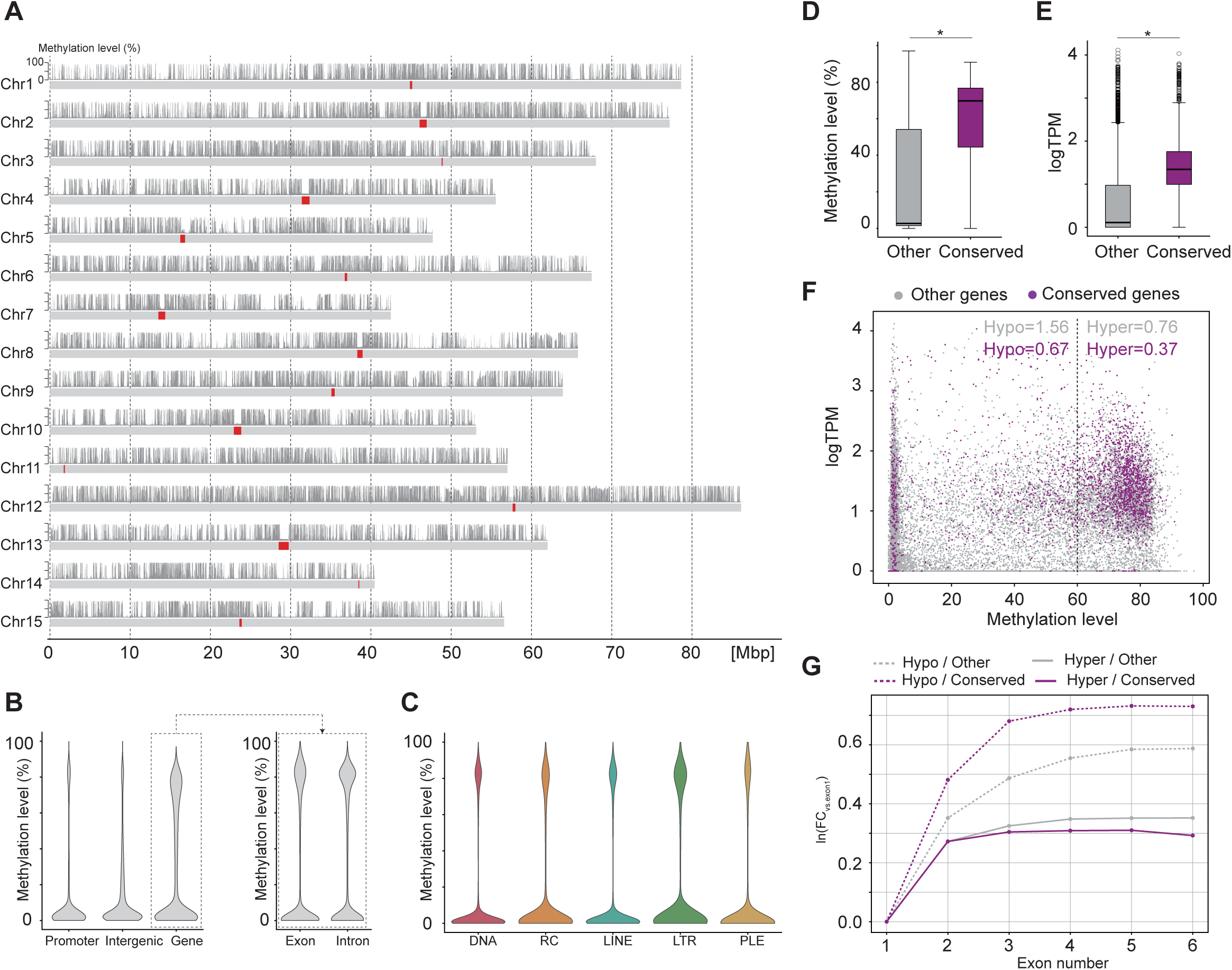
DNA Methylation within Gene Bodies Contributes to Transcriptional Stability (Homeostasis) of Conserved Genes in *H. vulgaris*. **(A)** Global pattern of DNA methylation in *H. vulgaris* AEP.JNIG genome showed that centromeric regions had low DNA methylation. The top track shows the methylation frequency (the probability that each base is methylated). The second track indicates the predicted centromere positions in red, which was identified by quarTeT (v 1.2.5) (Lin et al. 2023). **(B)** Methylation level across each genomic region. The gene region was further analyzed by dividing exon and intron regions. **(C)** Methylation level of each repetitive region. The colors correspond to those used in Fig 3A. **(D)** Boxplots of gene-body methylation levels in conserved (purple) and other (gray) genes. Differences were significant (Mann-Whitney U test, p < 1×10⁻⁴). **(E)** Boxplots showing expression levels in the conserved (purple) and the other (gray) genes. Differences were significant (Mann-Whitney U test, p < 1×10⁻⁴). **(F)** Relationship between DNA methylation levels in gene bodies and gene expression. Purple dots indicate conserved genes, and gray dots indicate other genes. Values indicate the coefficient variation of each group’s logTPM. **(G)** Spurious transcription in gene bodies is significantly lower in the conserved and highly methylated genes (normalized by exon length for the logFC). The y axis shows the natural logarithm of the coverage fold change (FC) of exons 1 to 6 versus exon 1, with lower values meaning less spurious transcription. **Alt text**: Composite figure with panels A–G illustrating that gene-body DNA methylation is associated with conserved genes and contributes to transcriptional stability in *Hydra vulgaris*. A is a genome-wide visualization of DNA methylation frequency showing low methylation levels in centromeric regions, with predicted centromere positions indicated along the chromosomes. B is a comparison of methylation levels across genomic regions, showing gene-body methylation patterns in exons and introns. C is a comparison of methylation levels across different classes of repetitive elements. D is a boxplot comparing gene-body methylation levels in conserved genes (purple) and other genes (gray), showing significantly higher methylation in conserved genes. E is a boxplot comparing gene expression levels between conserved and other genes, with conserved genes showing higher expression. F is a scatter plot showing the relationship between gene-body methylation and gene expression variability, with conserved genes (purple) showing more stable expression patterns. G is a plot measuring spurious transcription within gene bodies, indicating that conserved and highly methylated genes show lower levels of spurious transcription.

We next compared methylation levels across genomic features. DNA methylation levels in promoters, exons, introns, and intergenic regions exhibited bimodal distributions, with distinct peaks corresponding to highly and weakly methylated regions. The methylation levels within gene regions were higher on average than those within promoter and intergenic regions (Fig 3B and S4A Fig). Repetitive elements also exhibited a bimodal methylation pattern. Among repeat classes, LINEs and LTRs tended to be more highly methylated, a trend that is clearer in the boxplots (Fig 3C and S4B Fig).

We further analyzed the relationship between DNA methylation and gene expression in our dataset. We focused on conserved genes, defined as orthologs shared across the 33 holozoans (see Materials and Methods), because conserved genes have been reported to be hypermethylated in multiple species, including cnidarians (Zhang & Jacobs 2022; Sarda et al. 2012). Consistent with these reports, conserved genes in our data showed higher methylation than other genes (Fig 3D). Conserved genes tended to be more highly expressed and exhibited less variation in expression levels (Fig 3E and 3F), suggesting a link between methylation and stable gene regulation. Finally, the ratio of transcription levels between exon 1 and downstream exons, a measure of potential spurious transcription (Li et al. 2018), was negatively correlated with exon methylation, and this correlation was more pronounced for conserved genes than for other genes (Fig 3G).

These results suggest that genomic methylation, particularly within gene bodies, may help stabilize higher expression levels and suppress spurious transcription, especially in conserved genes. This observation is consistent with findings in other cnidarians, such as coral (Li et al. 2018).

### Aging-related orthologs are conserved in *Hydra* comparable to other aging model species

We then investigated aging-related orthologs in immortal *H. vulgaris* strains (105 (Chapman et al. 2010), AEP (Cazet et al. 2023), and AEP.JNIG (this study)) and the aging species *H. oligactis* (Cazet et al. 2023) in comparison with other model organisms, including *Homo sapiens*, *Mus musculus*, *Drosophila melanogaster*, and *Caenorhabditis elegans*. To reduce the impact of incomplete genome assemblies or gene annotations, we integrated orthogroup information across three *H. vulgaris* strains, treating an orthogroup as present when it was detected in any one of the strains.

First, we manually compiled a list of aging-related genes based on the *Hallmarks of Aging* framework (López-Otín et al. 2023). Specifically, genes explicitly mentioned in the original paper were directly included. For hallmarks described at the level of signaling pathways rather than individual genes, constituent genes were identified primarily using the KEGG Pathway database (Kanehisa & Goto 2000). When pathways were not clearly defined or fully represented in KEGG, additional genes were identified using the Gene Ontology (GO) database and relevant literature. The complete list of genes is summarized in S1 Table. Subsequently, we used OrthoFinder (Emms & Kelly 2019) to construct orthogroups based on predicted protein sequences derived from genome assemblies, supplementing the analysis with BLAST searches. Finally, we examined the orthogroups annotated with these aging-related genes. The presence of a sequence from a given species within an orthogroup was taken as evidence that the species possesses the corresponding gene.

Our analysis revealed that *Hydra* (both *H. vulgaris* and *H. oligactis*) conserve a substantial proportion of genes associated with key aging-related pathways, such as autophagy, proteostasis, DNA damage repair, mitochondrial quality control, stem cell regulation, insulin receptor signaling, NAD signaling, and cGAS-STING signaling (Fig 4A and S5 Fig). For instance, approximately 93% of the genes in the KEGG autophagy pathway (107/115) and 93% in the base excision repair pathway (39/42) were identified in *H. vulgaris* (see S2 Table for details). These findings are consistent with previous reports (Pascual-Torner et al. 2022; Barve et al. 2021; Chera et al. 2009) and suggest that the fundamental mechanisms of aging mediated by these genes are likely conserved across species, including the immortal *H. vulgaris* (Fig 4A).

**Figure 4.**
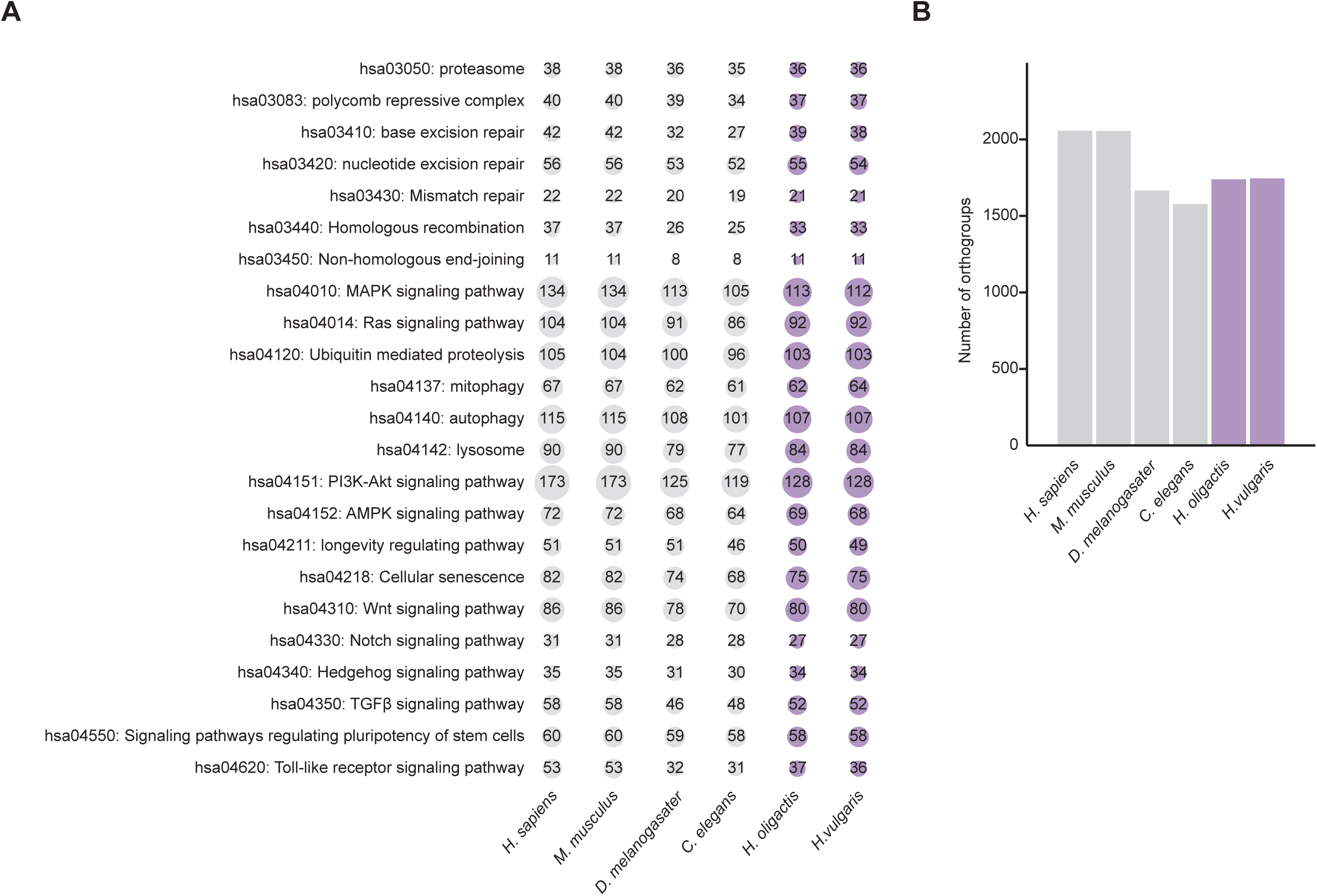
Conservation of aging-related orthologs in *Hydra*. **(A)** Conservation of orthologs related to the *Hallmarks of Aging*. The vertical axis lists representative aging-related pathways (defined as KEGG terms). Each pathway was extracted from the 12 hallmarks of aging proposed in 2023 (López-Otín et al. 2023). Genes including each pathway were listed as described in the Materials and Methods and summarized in S4 Table. Numbers inside the circles indicate the number of conserved orthologs, and the circle size corresponds to the number of conserved orthologs. The *Hydra* genus is highlighted in purple. **(B)** Bar plot showing the total number of conserved aging-related orthologs identified in each species. The y-axis represents the number of orthogroups. The *Hydra* genus is highlighted in purple. **Alt text**: Composite figure with panels A–B illustrating the conservation of aging-related genes across species. A is a bubble plot showing the conservation of orthologs associated with aging-related pathways defined by the Hallmarks of Aging. Pathways are listed on the vertical axis, and circle size indicates the number of conserved orthologs. The *Hydra* genus is highlighted in purple. B is a bar plot comparing the total number of conserved aging-related orthologs identified in each species, with *Hydra* species highlighted in purple, showing *Hydra* species have more aging-related orthologs than flies or nematodes.

Furthermore, a cross-species comparison of the total number of aging-related orthologs revealed that *Hydra* shares a larger number of conserved orthologs with humans than do *D. melanogaster* and *C. elegans* (Fig 4B). This indicates that *Hydra* has retained a highly conserved repertoire of aging-related genes, despite its significant evolutionary distance from traditional model organisms. Our results are supported by previous work using Reciprocal Best Hit (RBH) analysis, which similarly showed that *H. vulgaris* possesses a greater number of human orthologs than either *D. melanogaster* or *C. elegans* (Wenger & Galliot 2013).

Therefore, these results substantiate the potential of *Hydra* as a model organism for aging research, offering a unique opportunity to investigate conserved mechanisms of organismal aging across the metazoan tree of life.

### Comparative Analysis between *H. vulgaris* and *H. oligactis* reveals aging-related orthologs

Finally, we conducted a comparative orthogroup analysis between *H. vulgaris* and *H. oligactis*, particularly aiming at the identification of gene sets related to their aging or immortal traits. As was in Fig 4, we used the integrated dataset for *H. vulgaris* orthogroups. To confirm the absence of these orthogroups in each species, a BLAST search (Altschul et al. 1990) was performed (see Materials and Methods). This analysis revealed that *H. vulgaris* possessed 33 unique orthogroups that were not detected in *H. oligactis*, while *H. oligactis* contained 65 unique orthogroups (Fig 5A and 5B; S3 Table).

**Figure 5.**
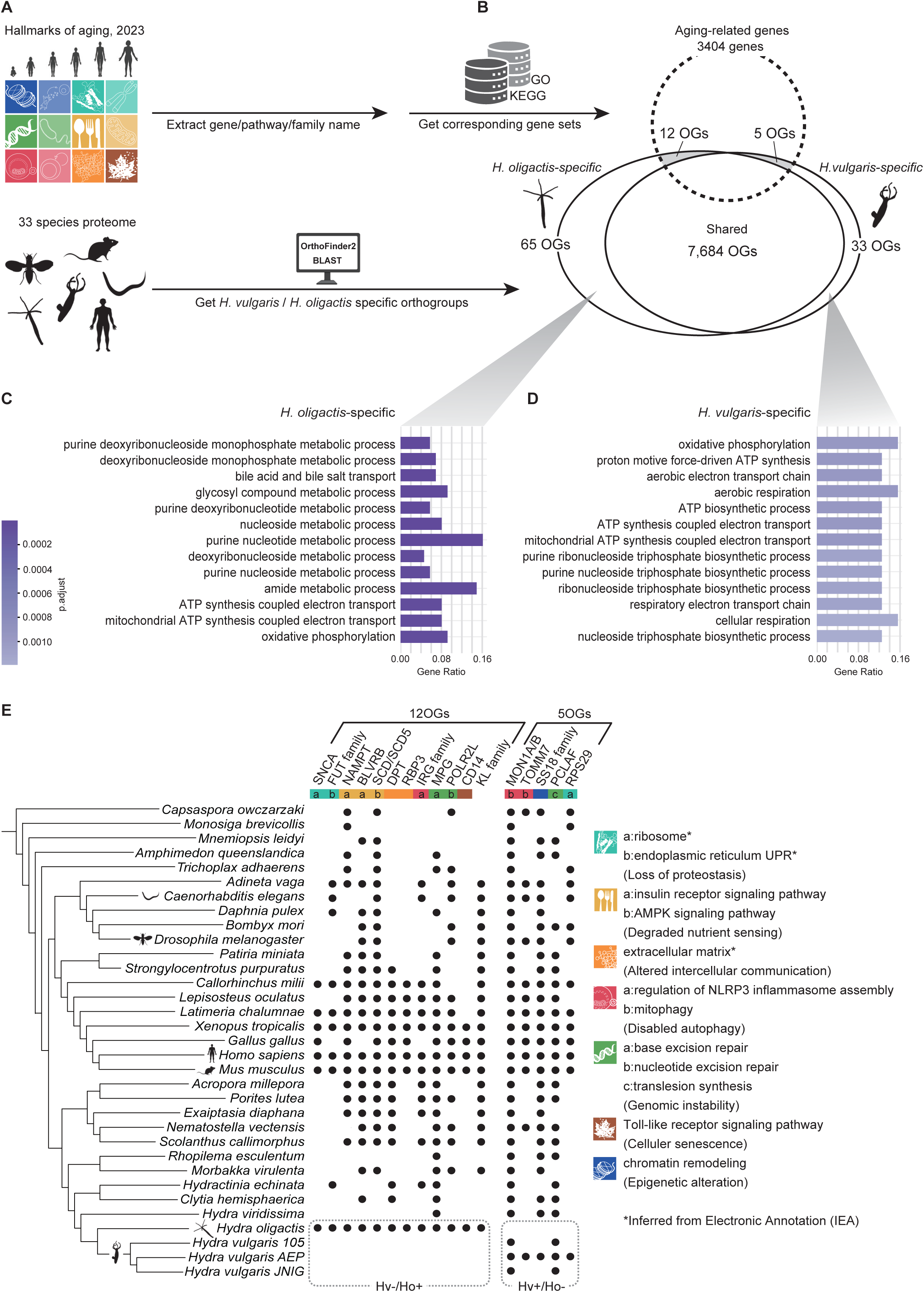
Comparative Analysis between *H. vulgaris* and *H. oligactis*. **(A)** Schematic diagram illustrating the method used to identify *H. vulgaris*- and *H. oligactis*-specific aging-related genes. A part of figure materials was prepared using biorender.com (permission No. JK2989SSUB). **(B)** Twelve genes are present in *H. oligactis* but absent in *H. vulgaris*, whereas five genes are present in *H. vulgaris* but not in *H. oligactis*. A total of 7,684 genes is shared between the two species. Note that only genes annotated based on homology to human genes were included in this analysis. We referred to the original article (López-Otín et al. 2023) for the factors of *Hallmarks of Aging*. Icons for the analyzed species were referred to from PhyloPic (https://www.phylopic.org/) under Creative Commons license. **(C)** Gene Ontology (GO) enrichment analysis for biological processes was performed using 65 orthogroups present in *H. oligactis* but absent in *H. vulgaris*. The y-axis shows the top 13 significantly enriched GO terms. The x-axis represents the gene ratio, defined as the proportion of input genes associated with each term. The color intensity of the bars indicates the adjusted p-value (p.adj), with darker colors representing higher statistical significance. **(D)** The same analysis was performed for 33 orthogroups present in *H. vulgaris* but absent in *H. oligactis*. **(E)** Orthogroups specific to *H. oligactis* (65 OGs) and *H. vulgaris* (33 OGs) were screened across 33 organisms to identify those involved in aging-related pathways defined in this study based on the *Hallmarks of Aging*. The final list of genes is shown on the x-axis. The y-axis represents the species included in this analysis, ordered by phylogeny. Dots indicate the presence of the corresponding gene in each organism, while blanks indicate absence. The colors at the top represent the *Hallmarks of Aging* categories assigned to each gene. The names of each category and the specific terms included in them are shown on the right. **Alt text**: Composite figure with panels A–E illustrating comparative analyses of aging-related genes between *Hydra vulgaris* and *Hydra oligactis*. A is a schematic diagram showing the analytical workflow used to identify species-specific aging-related genes in the two *Hydra* species. B is a diagram summarizing shared and species-specific genes, showing 7,684 genes shared between the species, 65 genes present only in *H. oligactis* (12 of them aging-related), and 33 genes present only in *H. vulgaris* (5 of them aging-related). C is a bar plot showing Gene Ontology enrichment results for orthogroups present in *H. oligactis* but absent in *H. vulgaris*, with the y-axis listing enriched biological processes and the x-axis showing the gene ratio. D is a similar Gene Ontology enrichment plot for orthogroups present in *H. vulgaris* but absent in *H. oligactis*. E is a presence–absence matrix showing the distribution of species-specific orthogroups across 33 organisms, with dots indicating gene presence and blank cells indicating absence.

To explore their biological functions, we performed Gene Ontology (GO) enrichment, KEGG pathway, and Reactome pathway analyses using clusterProfiler (v4.16.0) (Wu et al. 2021) (Figs 5C and 5D; S6 Fig; S4 Table). Intriguingly, terms related to oxidative phosphorylation (OXPHOS) were enriched in both species (e.g., adjusted p-value = 4.60E-5 for *H. oligactis* and 9.72E-4 for *H. vulgaris* in GO-biological processes (GO-BP); S4 Table). However, the specific genes contributing to these terms are contrasted. Unique genes identified in the *H. oligactis* GO-BP terms included α-synuclein (SNCA), mitochondrial deoxyguanosine kinase (DGUOK), and components of complexes I (NDUFA10, MT-ND3), III (MT-CYB), IV (MT-CO2/CO3), and V (MT-ATP6). In contrast, *H. vulgaris* uniquely possessed different components of complexes I (NDUFA3/B7), III (UQCR10/11), and V (ATP5F1E). The analysis also revealed species-specific terms regarding nucleic acid metabolic processes. For instance, *H. oligactis*-specific orthogroups were enriched for GO-BP terms such as “purine deoxyribonucleoside monophosphate metabolic process” (adjusted p-value = 8.29E-7), “purine nucleotide metabolic process” (adjusted p-value = 2.99E-6), and “amide metabolic process” (adjusted p-value = 3.71E-5) (Fig 5C and S4 Table). Conversely, in *H. vulgaris*, major GO-BP terms related to NTP synthesis were enriched, such as “ATP biosynthetic process” (adjusted p-value = 9.72E-4) and “purine nucleoside triphosphate biosynthetic process” (adjusted p-value = 1.10E-3) (Fig 5D and S4 Table). These contrasting results suggest species-specific characteristics in energy production and metabolic processes.

We further narrowed our analysis to aging-related genes and pathways unique to each species. By intersecting the species-specific orthogroups with the aging-related genes associated with the *Hallmarks of Aging* (Fig 4 and S1 Table), we identified twelve [SNCA, FUT1/2, NAMPT, BLVRB, SCD/SCD5, DPT, RBP3, IRG family (IRGC, IRGM), MPG, POLR2L, CD14, Klotho family (KLB, LCT, LCTL, KL, GBA3)] and five [MON1A/1B, TOMM7, SS18 family (SS18, SS18L1/2), PCLAF, RPS29] unique orthogroups for *H. oligactis* and *H. vulgaris*, respectively (Fig 5B). Extending our comparative analysis to 29 other holozoan species revealed that the twelve aging-related orthogroups identified in *H. oligactis* are largely conserved across these taxa. Notably, both humans and mice possess the complete set of these twelve orthogroups (Fig 5E). Conversely, the five orthogroups initially identified as unique to the immortal *H. vulgaris* were found to be present in the other 29 species (Fig 5E). This suggests that these orthogroups are unlikely to be the primary drivers of the immortal phenotype. Consequently, we focused our subsequent investigation on the twelve *H. oligactis*-specific orthogroups.

These twelve orthogroups possess well-known functions in aging processes. α-synuclein (SNCA) is a key molecule implicated in Parkinson’s disease and mitochondrial oxidative phosphorylation (OXPHOS) deficiency (Miwa et al. 2022). Nicotinamide adenine dinucleotide (NAD) is widely recognized for its association with aging (Imai & Guarente 2014; Imai 2016), and upregulation of genes involved in NAD^+^ biosynthesis accompanies lifespan extension (Tyshkovskiy et al. 2023). Nicotinamide phosphoribosyltransferase (NAMPT) functions as a rate-limiting enzyme in NAD^+^ biosynthesis, while biliverdin reductase B (BLVRB) is an NAD(P)H-dependent reductase involved in antioxidant defense. Stearoyl-CoA desaturase (SCD/SCD5) has been linked to brain aging in humans and mouse models of Alzheimer’s disease through its role in the synthesis of unsaturated fatty acids (McNamara et al. 2008; Hamilton et al. 2022). N-methylpurine DNA glycosylase (MPG) is involved in the base excision repair (BER) pathway, potentially contributing to the DNA damage response associated with aging, and is known to be upregulated in long-lived mammals (Tyshkovskiy et al. 2023; Xu et al. 2008). The presence of RNA polymerase II, I, and III subunit L (POLR2L) may be related to aging-associated changes in transcriptional integrity (Debès et al. 2023; Papantonis et al. 2025). Finally, it is intriguing that Klotho family molecules are absent in *H. vulgaris*. *Klotho* is a well-known anti-aging gene whose knockout in mice causes premature aging (Kuro-o 2009). Although the absence of Klotho in the immortal *H. vulgaris* appears inconsistent with the findings in mice, it suggests that the dependence on Klotho-mediated pathways, such as phosphate and vitamin D metabolism, may be essential for conferring the aging traits.

Collectively, these comparative analyses offer insights to address fundamental biological questions: specifically, how *H. vulgaris* acquired the immortal trait (or conversely, lost the aging trait) and what constitutes the essential genes and pathways underlying organismal aging.

## Discussion

The purpose of this study was to establish a chromosome-level genome assembly and accompanying epigenomic resources for *H. vulgaris* strain AEP.JNIG and offer insights into *H. vulgaris* immortality through comparative genome analysis.

We established a chromosome-level genome assembly for *H. vulgaris* AEP.JNIG with quality metrics comparable to the previously reported AEP strain (BUSCO: 94.7%; Scaffold N50: 63.9 Mb). Using the new high-quality genome assembly and accompanying gene annotations, we clarified the relationship of our strain to other *H. vulgaris* strains. Our analyses demonstrate that *H. vulgaris* AEP.JNIG forms a monophyletic group with *H. vulgaris* AEP. This close relationship was consistently supported by both a phylogenomic dataset including multiple *H. vulgaris* strains as well as single-gene analyses incorporating a broader taxonomic sampling of *Hydra* laboratory strains and natural isolates.

Despite the phylogenetic proximity of the *H. vulgaris* AEP.JNIG and *H. vulgaris* AEP, through detailed comparison between the two genomes, we detected a substantial accumulation of SNPs and indels, along with numerous inversions and duplications. These genomic variations likely reflect “laboratory drift” driven by long-term asexual reproduction, consistent with recent findings that asexual lineages accumulate somatic mutations at high rates (Kershenbaum et al. 2025; Sahm et al. 2024) and undergo structural changes such as Robertsonian translocations (Kon-Nanjo et al. 2025). Together, these findings highlight substantial genomic diversity within the species, establishing the AEP.JNIG assembly as a crucial resource that complements the existing *H. vulgaris* genomes.

In addition to the genome assembly, we provide foundational resources for epigenomic analyses, including single-nucleotide 5-methylcytosine (5mC) calls and transposable element (TE) annotations. Our Nanopore signal-based analysis identified a global methylation level of 19%, which is lower than the 27% previously estimated by bisulfite sequencing (Ying et al. 2022). This difference likely reflects the improved accuracy of signal-based detection, which is particularly advantageous for analyzing repeat-rich genomes like *Hydra* (Doshi et al. 2025). We confirmed that the *H. vulgaris* genome exhibits a characteristic methylation feature of invertebrates (Suzuki & Bird 2008), where gene bodies are hypermethylated while promoters and putative centromeres remain hypomethylated. Further, we found the potential functional link between DNA methylation and transcriptional stability. We observed that hypermethylation within gene bodies, particularly in evolutionary conserved genes, correlates with high expression levels and low transcriptional noise (spurious transcription). This finding suggests that gene body methylation serves as a mechanism to maintain transcriptional fidelity and homeostasis in *Hydra*, aligning with findings in other cnidarians such as corals (Li et al. 2018; Dixon et al. 2018). The robust methylation-based stabilization of the transcriptome, together with the loss of an RNA polymerase component (POLR2L) (Fig 5), may represent a key machinery for *Hydra*’s negligible senescence. Given that epigenetic alterations relates to the progress of aging, and that species-specific epigenetic signatures have recently been shown to determine maximum lifespan in mammals (Li et al. 2024), *Hydra* serves as a unique model for investigating how an organism can escape these epigenetic constraints.

Finally, to interrogate the genetic basis of *Hydra* immortality, we employed a comparative genomics approach contrasting the immortal *H. vulgaris* with the aging-inducible *H. oligactis*, consistent with recent frameworks emphasizing comparisons between exceptional species and related controls (Rechsteiner et al. 2025). Our orthology analysis confirmed that *Hydra* shares a larger repertoire of aging-related orthologs with humans than do traditional invertebrate models (Wenger & Galliot 2013) and highly conserves genes associated with the *Hallmarks of Aging*. This supports the view that aging mechanisms are ancestral (Lemoine 2021), suggesting *Hydra* achieves immortality by robustly regulating these pathways rather than lacking them.

Intriguingly, however, we identified a counter-intuitive divergence: potential anti-aging genes (e.g., Klotho, NAMPT, MPG) are unique to the aging *H. oligactis* and absent in the immortal *H. vulgaris*. Additionally, metabolic signatures differed, with *H. vulgaris* enriched for ATP synthesis pathways and *H. oligactis* for amide/purine metabolism pathways. We hypothesize that *H. vulgaris*, inhabiting thermally stable environments (Khalturin et al. 2009), has optimized an efficient energy metabolism that minimizes cellular damage, such as the synthesis of reactive oxygen species. In contrast, the presence of metabolic regulators like Klotho in *H. oligactis*, living in a cooler environment and sensitive to heat (Khalturin et al. 2009; Bosch et al. 1988), implies complexity and trade-offs in the metabolic system, consistent with an evolutionary aging theories such as antagonistic pleiotropy (Kirkwood 1977; Hamilton 1966). Thus, *H. vulgaris* may maintain immortality by relying on a simpler homeostatic system that avoids dependence on signaling pathways prone to age-dependent deterioration and damage accumulation. From the viewpoint of evolutionary aging theories, it would be intriguing to investigate mechanisms by which sexual reproduction triggers senescence in *H. oligactis*, particularly within the context of the metabolic trade-offs imposed by its distinct genetic background.

Several limitations of this study should be noted. First, our comparative analysis relies on orthology with human aging-related genes defined by the *Hallmarks of Aging*. This approach may overlook lineage-specific aging mechanisms unique to cnidarians. Additionally, despite rigorous BLAST validation (e-value < 10^-5^), the large evolutionary distance may lead to potential undetected orthologs due to sequence divergence. Second, and most importantly, the functional roles of the identified candidate genes (e.g., *H. oligactis*-specific genes) and the causal link between DNA methylation and transcriptional stability remain to be experimentally verified. Future studies employing functional assays, such as gene knockdown or transgenic approaches, are essential to confirm the biological significance of these genomic findings for *Hydra* aging and immortal traits.

In conclusion, our study provides an additional genomic resource comprising a new high-quality, chromosome-level assembly of *H. vulgaris* AEP.JNIG, which facilitates the analysis of genomic variation across independent laboratory populations. We also present a robust DNA methylation profile derived from Nanopore sequencing signals, offering a foundation for future epigenetic studies in cnidarians. Furthermore, through comparative genomics, we offer novel insights into the potential genomic origins of *H. vulgaris* immortality, highlighting distinct metabolic strategies and the evolutionary trade-offs associated with anti-aging genes. Collectively, these resources and findings further establish *Hydra* as a robust and unique model for advancing the comparative biology of aging. To fully interrogate the questions regarding immortality in *Hydra*, more extensive analyses are required, including investigations into individual gene functions, the system-level input-output characteristics of entire pathways, and genetic loss- and gain-of-function assays.

## Materials and Methods

### *Hydra* strain and culture

A *Hydra vulgaris* strain maintained as AEP at the Japan National Institute of Genetics (JNIG) was acquired and cultured at Juntendo University using standard procedures. Briefly, the animals were kept at 20°C in the *Hydra* medium (1 mM Tris-HCl pH7.6, 1 mM CaCl2, 1 mM NaCl, 0.1 mM KCl, 0.1 mM MgSO4) and were fed brine shrimp (*Artemia franciscana*) three times a week.

### Genome and RNA-Seq sequencing

For clean genomic DNA extraction, 100 polyps of *H. vulgaris* AEP.JNIG polyps were starved for 10 days in the presence of antibiotics (50 μg/mL of ampicillin, rifampicin, neomycin, and streptomycin) and herbicide (3-(3,4-dichlorophenyl)-1,1-dimethylurea (DCMU), 5 μM) to eliminate possible algal contaminants. Whole genomic DNA was then extracted using WiZard HMW DNA Extraction Kit (Promega, Cat. No. A2920) and sent for sequencing.

Nanopore sequencing was done using the instrument PromethION with the flowcell version R.9.4.1. The basecalling was done using Guppy version 5.0 (https://nanoporetech.com/ja/software/other/guppy/). The qualities of the reads were assessed using NanoPlot (https://github.com/wdecoster/NanoPlot) (De Coster & Rademakers 2023). Short-read genome sequencing libraries were also prepared from the extracted genomic DNA using the Truseq DNA PCR-Free kit and 150 bp paired-end reads were sequenced on an Illumina NovaSeq 6000 platform.

The Hi-C sequencing library was prepared from 100 polyps of *H. vulgaris* AEP.JNIG with EpiTect Hi-C Kit (Qiagen) according to manufacturer’s instructions and was sequenced on NovaSeq X Plus (Illumina) as 150 bp paired-end reads.

### Karyotyping

Karyotyping was performed with minor modifications to a previously described protocol for various invertebrates (Guo et al. 2018). To increase the number of proliferating cells (David & Campbell 1972), *H. vulgaris* AEP.JNIG polyps were heavily fed in the morning and, 12 hours later, treated with 0.1% nocodazole for 1 hour at room temperature. The organisms were then fixed in Carnoy’s fixative (methanol : acetic acid = 3 : 1) for 30 minutes on ice. Fixed *Hydra* bodies were transferred to glass slides and incubated with a few drops (∼30 µL) of 60% acetic acid for 5 minutes at room temperature. Siliconized coverslips prepared with Sigmacote (Sigma-Aldrich, Cat. No. SL2-25ML) were placed on top of the specimens, and the tissues were gently squashed by applying constant pressure for approximately 3 seconds to obtain a single layer of nuclei (Guo et al. 2018). Slides were incubated overnight at 4 °C and subsequently chilled on dry ice for 10 minutes. Coverslips were removed using a blade while the slides remained on dry ice. After returning to room temperature, slides were gently rinsed twice with 1x phosphate-buffered saline (PBS), stained with 1 µg/mL 4′,6-diamidino-2-phenylindole (DAPI) for 5 minutes at room temperature, rinsed again with PBS, and mounted using ProLong Diamond (Invitrogen, Cat. No. P36965). Samples were imaged using an in-house spinning-disk confocal microscope (Nikon Eclipse Ti2-E equipped with a spinning-disk scanner with 100-μm-wide pinholes aligned with a Nipkow disk (CSU-MPϕ100; Yokogawa Electric) (Otomo et al. 2015) and an oil-immersion lens (SR HP Apo TIRF 100XC Oil, numerical aperture [NA] = 1.49, Nikon)).

### Genome assembly and annotation

The fastq files were assembled using the Phased Error Correction and Assembly Tool (PECAT) (Nie et al. 2024) with medaka (https://github.com/nanoporetech/medaka) as a polishing tool in the final phase of assembly.

For Hi-C scaffolding in *H. vulgaris* AEP.JNIG basecalled and demultiplexed reads totaling 119 Gbp were first filtered using fastp v0.19.4 (Chen 2023) with -c and -g options, and the filtered reads were processed with HiC Pro v3.1.0 (Servant et al. 2015). Qualified reads were used to scaffold the assembly using YAHS v1.2 (Zhou et al. 2022) with default parameters. Scaffolds with length over 10 kbp were retained. Contact map was generated using Juicer 2.20 pipeline (Durand et al. 2016). The Hi-C contact map was curated using the Juicebox v1.11.08 (Durand et al. 2016). The final Hi-C contact map and genome assembly were generated using the ‘run-asm-pipeline-post-review.sh’ script from the 3D-DNA pipeline (Dudchenko et al. 2017). The completeness of filtered genome assembly was evaluated by BUSCO v5.7.1 (Manni et al. 2021) with the metazoa_odb10 dataset.

Prior to gene annotation, we soft-masked the assembled genome using RepeatModeler v2.0.2 and RepeatMasker v4.1.1 (Flynn et al. 2020). The soft-masked genomes were used to annotate protein-coding genes using BRAKER v2.1.6 (Brůna et al. 2021). We used *H. vulgaris* AEP.JNIG whole-body RNA-seq datasets as hints for annotation, mapped with HISAT2 v.2.2.1 (Kim et al. 2019).

The gene models annotated for the initial genome assembly were transferred to the final assembly using Liftoff v1.6.3 (Shumate & Salzberg 2021). Nine transcripts that were not transferred properly with the initial Liftoff run were manually mapped to the new assembly using minimap2 (Li 2018) with the splice mode, and the ORFs were predicted using Transdecoder (Haas).

### Phylogenetic tree reconstruction

We constructed a proteome dataset of 31 metazoan and 2 unicellular holozoan organisms. A workflow of constructing multiple sequence alignments based on the one described in (Liu et al. 2024) was followed. Briefly, BUSCO v.5.8.0 (Manni et al. 2021) (Manni et al. 2021)was performed against each proteome using metazoa_odb10. Single-copy genes were extracted from the BUSCO output of each species and collected into individual FASTA files. These FASTA files were filtered first by the number of species present in the dataset, with the threshold being 23 (i.e., less than 10 missing species). PREQUAL v1.02 (Whelan et al. 2018) was run with a posterior probability threshold of 0.95. Then MAFFT v7.525 (Katoh & Standley 2013) was used with the “-linsi” mode, without any other optional parameters. Divvier v1.01 (Ali et al. 2019) was used to remove regions of uncertain alignment (with the “-mincol 4 - divvygap” option), followed by trimming using trimAl v1.5 (Capella-Gutiérrez et al. 2009) (with the “-automated1” option). All the individual alignments were concatenated to create the final supermatrix used for the subsequent phylogenetic analyses. A maximum-likelihood phylogenetic tree was reconstructed using IQ-TREE2 v2.3.6 (Minh et al. 2020) under the “LG+C60+F+G” model, with branch support assessed by the ultrafast bootstrap method (1,000 replicates).

Molecular phylogenetic trees of *Hydra* strains based on mitochondrial and nuclear genes (*COI*, *COII*, *EF1α*, and *Cnnos1*) were reconstructed using the dataset used in (Kawaida et al. 2010). Corresponding genes were retrieved from the *H. vulgaris* AEP.JNIG and the *H. vulgaris* AEP genomes and added to the original dataset. All trees were reconstructed with IQ-TREE2 under the best-fitting substitution models selected by ModelFinder (MFP option), with branch support assessed using the ultrafast bootstrap method.

### Detection and visualization of genomic rearrangements between *H. vulgaris* AEP.JNIG and AEP assemblies

A dot plot was generated with D-GENIES (Cabanettes & Klopp 2018) using the genomes of *H. vulgaris* AEP.JNIG and *H. vulgaris* AEP. Gene synteny plot was created using GENESPACE (Lovell et al. 2022). Structural rearrangements between the two assemblies were then detected by SyRI v1.7.1 (Goel et al. 2019) using *H. vulgaris* AEP as the reference and *H. vulgaris* AEP.JNIG as the query, and the results were visualized with plotsr (Goel & Schneeberger 2022). Gene density was calculated based on *H. vulgaris* AEP gene models. Centromere positions were estimated from the repeat annotations identified by EDTA v2.2.2 (Ou et al. 2019) and processed with quarTeT (v1.2.5) (Lin et al. 2023), while repeat density was calculated using the output from Earl Grey (v6.3.3) (Baril et al. 2024).

### Transposable element prediction

Transposable elements (TEs) in the *H. vulgaris* AEP and AEP. JNIG genomes were identified using Earl Grey v6.3.3 (Baril et al. 2024).

### Methylation landscape analysis

The distribution of DNA methylation (mCpG) was detected from Nanopore raw signal data using Dorado (v0.7.1) (Oxford Nanopore Technologies: Dorado. https://github.com/nanoporetech/dorado.) and Modkit (v0.4.3) (Oxford Nanopore Technologies: Modkit. https://github.com/nanoporetech/modkit.). In Fig 3A the methylation frequency values output by Modkit were visualized using karyoploteR v1.32.0 (Gel & Serra 2017). Based on the gene model of *H. vulgaris* AEP.JNIG and the genomic locations of each TE class detected by Earl Grey v6.3.3 (Baril et al. 2024), mCpG/CpG ratios calculated for each genomic region using the “modkit stats” command in Fig 3B and 3C. Promoter regions were defined as the 2,000 bp upstream of the transcription start site (TSS), and regions that were neither within genes nor promoters were defined as intergenic regions. In Fig 3D and 3E, conserved genes were defined as orthologs shared across 33 species based on the output from Orthofinder. Fig 3F displays a scatter plot in which the x-axis represents the gene body–specific mCpG/CpG ratios obtained from “modkit stats”, and the y-axis shows the corresponding RNA-seq expression levels (logTPM). The conserved genes shown in Fig 3E correspond to orthologs shared among all 33 species analyzed in Fig 5.

### Spurious transcription analysis

Paired-end RNA-seq reads were aligned to the *H. vulgaris* AEP.JNIG genome using HISAT2 v2.2.1 (Kim et al. 2019) with default parameters. The resulting SAM files were converted to BAM format and sorted using SAMtools v1.23 (Danecek et al. 2021). Base-level read coverage across the genome was calculated using bedtools2 v2.31.1 (Quinlan & Hall 2010) with the command “bedtools genomecov -d”. Exon-level mean coverage was then calculated by mapping the exon BED file to the per-base coverage BED file using “bedtools map -o mean”. For each gene, exon-level coverage values were normalized by dividing by the coverage of the first exon (exon 1) of the same gene, ratios were log-transformed using the natural logarithm for downstream comparisons. Exons with zero or undefined coverage values were excluded. Genes were further annotated based on two criteria: DNA methylation status and evolutionary conservation. Methylated genes were defined as genes with mCpG/CpG ratios greater than 60%, whereas conserved genes were defined as genes belonging to orthogroups shared across all 33 species. This produced four groups: conserved–hypermethylated, conserved–hypomethylated, nonconserved–hypermethylated, and nonconserved–hypomethylated. For each exon position (exons 1–6), the mean log-transformed coverage ratio was calculated separately for each of the four gene groups.

### Conservation of aging related orthologous comparative Analysis of Orthologous Groups in *Hydra oligactis* and *Hydra vulgaris*

Orthologs were identified using OrthoFinder(v2.5.5) (Emms & Kelly 2019). Protein sequences from 33 species were used as input data, and the topology of the phylogenetic tree was based on the study by (DeBiasse et al. 2022; Schultz et al. 2023), which was then confirmed by our own phylogenomic analysis. Details are provided on the official GitHub repository (https://github.com/nojiruko/H_vulgaris_AEP.JNIG). Each orthogroup was annotated using human protein sequences as references, and the correspondence table of gene name and Protein stable ID obtained from Ensembl BioMart (https://www.ensembl.org/info/data/biomart/index.html) was used. Additionally, we compared the presence or absence of orthogroups between *Hydra oligactis* and *Hydra vulgaris, made a list of ortholog groups differentially present,* and performed GO, KEGG, and Reactome enrichment analyses. Enrichment analyses were conducted using ClusterProfiler v4.14.4 (Wu et al. 2021) and ReactomePA v1.50.0 (Yu & He 2016), with the human database referenced from <u>org.Hs.eg.db</u> v3.20.0 (Carlson et al. 2019).

Genes involved in aging-related pathways were determined based on López-Otín, C. *et al*. (2023) (López-Otín et al. 2023). First, we manually compiled a list of aging-related genes based on the *Hallmarks of Aging* framework (López-Otín et al. 2023). Specifically, genes explicitly mentioned in the original paper were directly included. For hallmarks described at the level of signaling pathways rather than individual genes, constituent genes were identified primarily using the KEGG Pathway database (Kanehisa & Goto 2000). When pathways were not clearly defined or fully represented in KEGG, additional genes were identified using the Gene Ontology (GO) database and relevant literature. The complete list of genes is summarized in S1 Table. For orthogroups showing differential conservation between *H. oligactis* and *H. vulgaris*, we performed BLASTP (protein query vs protein database) searches to verify whether homologous sequences were present in the proteome of the other species, using an e-value threshold of 10^-5^. Specifically, for orthogroups identified in *H. oligactis* but absent in *H. vulgaris*, *H. oligactis* protein sequences were used as BLASTP queries against the locally created proteome databases of *H. vulgaris* AEP and *H. vulgaris* AEP.JNIG. Conversely, for ortholog groups found in *H. vulgaris* but absent in *H.oligactis*, *H. vulgaris* AEP.JNIG protein sequences were queried against the locally created proteome database of *H. oligactis*. Orthogroups present in at least one *H. vulgaris* strain but not detected in *H. oligactis* were defined as *H. vulgaris*-unique. Orthogroups absent from all three *H. vulgaris* strains but present in *H. oligactis* were defined as *H. oligactis*-unique.

## Data Availability

The genome assembly of *H. vulgaris* AEP.JNIG and the associated sequence reads are available at GenBank under BioProject PRJNA1413991 (BioSample SAMN54890366). Raw Nanopore reads, Illumina genomic reads, and Illumina Hi-C reads have been deposited in the Sequence Read Archive (SRA) under accession numbers SRR37338222, SRR37338221, and SRR37338220, respectively.

## Acknowledgements

We thank the lab members at DBSB Juntendo, particularly Y. Kumagai, Q. Su, K. Suzuki, A. Kanemitsu, K. Matsumoto, Y. Saito, and M. Ota for maintaining *Hydra* polyps and supporting the related experiments; K. Otomo for supporting chromosome imaging and karyotyping; Y. Wada and T.K. Suzuki for supporting computational resources. We also thank GeneBay, Inc. (Yokohama, Japan) for sequencing the *H. vulgaris* AEP.JNIG genome; K. Ikeo (National Institute of Genetics, Japan) for maintaining and providing Japanese hydra resource at NIG, S. Kuraku (National Institute of Genetics, Japan), R. Nakato (The University of Tokyo, Tokyo, Japan), and A. Watanabe (CyberomiX Inc., Kyoto, Japan) for fruitful suggestions about genome assembly and analysis. We would also like to thank S. Hamada, K. Hamada, and O. Koizumi (Fukuoka Women’s University, Fukuoka, Japan) for his/her essential guidance in establishing *Hydra* in our laboratory and for providing protocols for its successful husbandry. We also extend our gratitude to Y. Takai (Institute for Advanced Biosciences, Keio University) for her technical assistance in Hi-C library preparation. We acknowledge the use of large language models (ChatGPT and Gemini) for English language proofreading and stylistic editing of the manuscript.

This work was supported by Japan Society for the Promotion of Science (JSPS) KAKENHI Grant-in-Aid for Scientific Research (B) [grant number JP22H02824 to E.A.S.]; JSPS KAKENHI for International Leading Research [grant number 23K20044 to E.A.S.]; Japan Science and Technology agency (JST) Core Research for Evolutionary Science and Technology (CREST) [grant number JPMJCR23B7 to E.A.S.]; Japan Agency for Medical Research and Development (AMED) Project for Promotion of Cancer Research and Therapeutic Evolution (P-PROMOTE) [grant number JP25ama221610 to E.A.S.]; Grants-in-Aid from Nakatani Foundation [to E.A.S.]; JSPS KAKENHI Gran-in-Aid for Research Activity Start-up [24K23213 to K.K.]; intramural Grant-in-Aid and Operating Costs Subsidies for Private Universities [to E.A.S.]; research funds from the Yamagata Prefectural Government and Tsuruoka City, Japan [to K.A.]; and the JSPS Overseas Research Fellowships [to T.K.].

## Author contributions

E.A.S., K.N., K.K., and A.S. conceived and directed the study. H.S. maintained and provided *H. vulgaris* AEP.JNIG resource. A.S. prepared the *H. vulgaris* AEP.JNIG genomic DNA and RNA samples for sequencing. K.A. performed the Hi-C sequencing and analysis. K.K., T.K., K.K-N., and K.A. assembled the genome data. K.N. and K.K. performed the genomic comparison between AEP.JNIG and AEP, whole-genome methylation analysis, and comparative genome analysis. K.K. performed the karyotyping experiment. E.A.S., T.K., and K.A. acquired funding. E.A.S., K.N., and K.K. mainly wrote the manuscript. All authors discussed the results and commented on the manuscript.

## Supplementary figure legends

**Figure S1.**
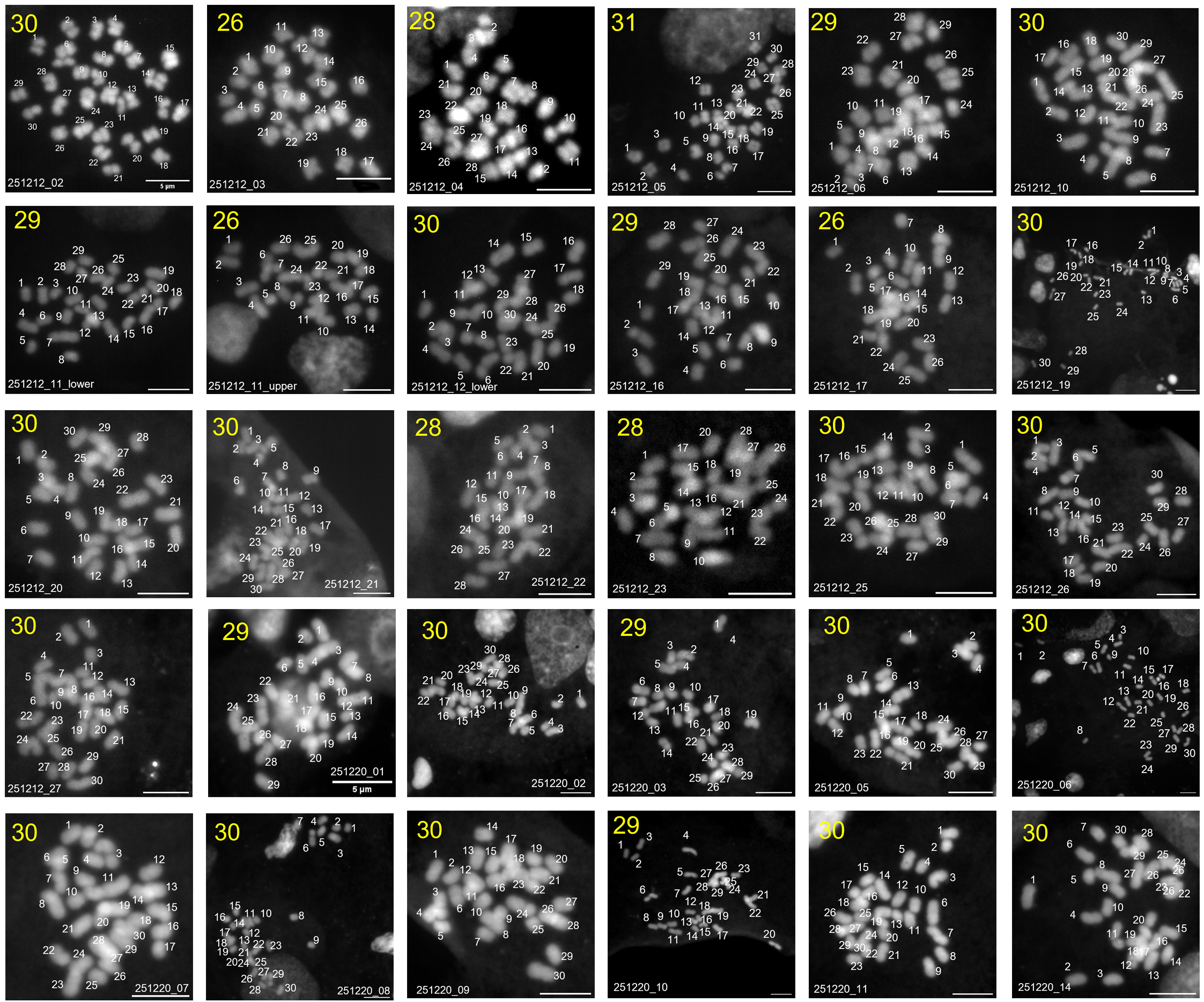
Karyotyping Data. Karyotype of *H. vulgaris* AEP.JNIG. A total of 30 nuclei samples were examined, and the number of chromosomes were counted. The number in yellow indicates the karyotype (2n) of the corresponding image. Scale Bar = 5 μm.

**Figure S2.**
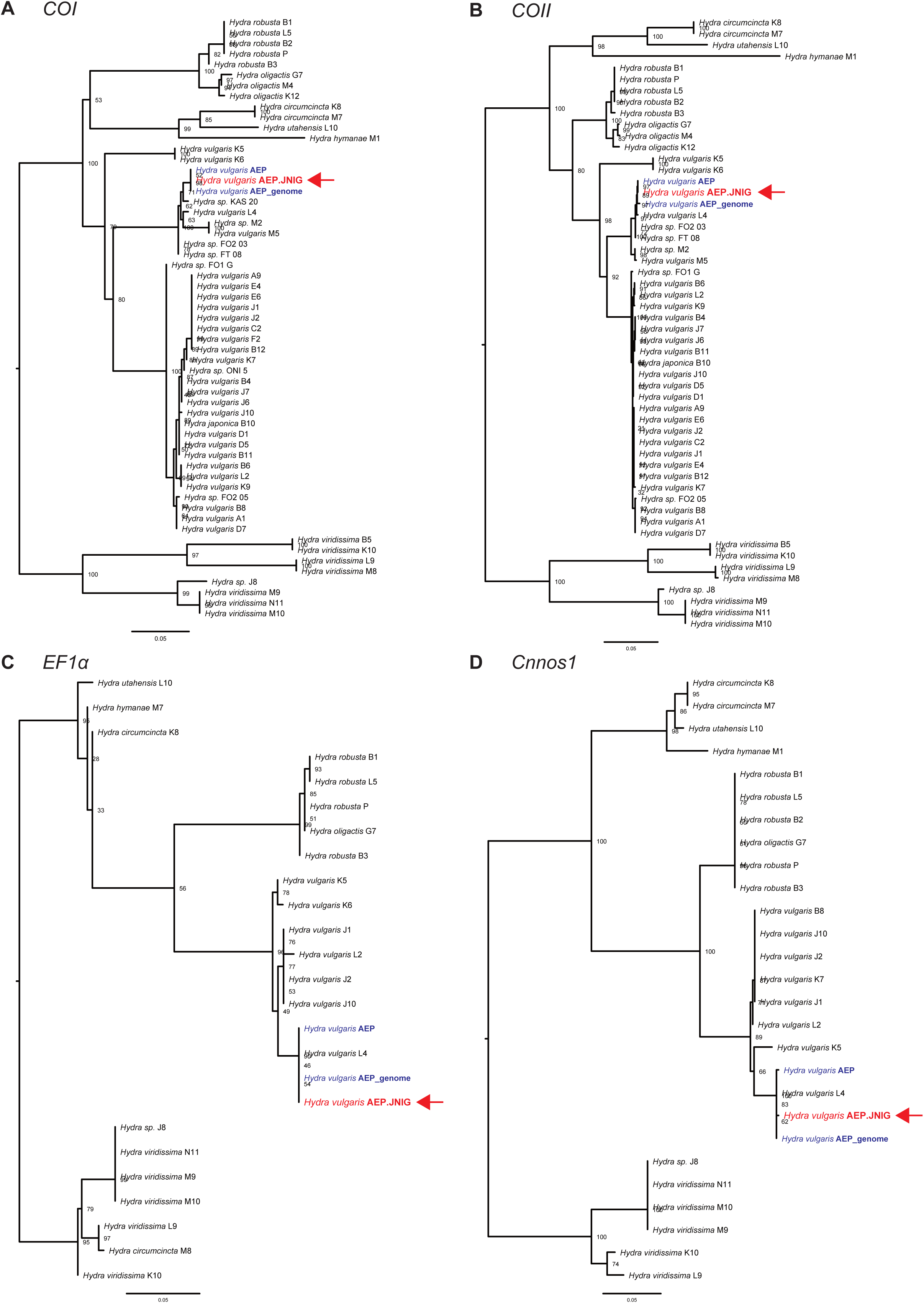
Molecular phylogenetic trees of *Hydra* strains based on mitochondrial and nuclear genes. Trees were reconstructed using IQ-TREE2 with best-fitting models selected by ModelFinder (MFP option). Branch support was evaluated using the ultrafast bootstrap method. *H. vulgaris* AEP.JNIG and *H. vulgaris* AEP sequences are shown in red and blue, respectively. *H. vulgaris* AEP sequences from (Kawaida et al. 2010) (“*Hydra vulgaris* AEP”) and those derived from the *H. vulgaris* AEP genome (Cazet et al. 2023) (“*Hydra vulgaris* AEP_genome”) (Cazet et al. 2023)are indicated by distinct labels. Numbers at nodes represent ultrafast bootstrap support values. Scale bars represent 0.05 nucleotide substitutions per site. **(A)** A molecular phylogenetic tree based on *COI*. **(B)** A molecular phylogenetic tree based on *COII*. **(C)** A molecular phylogenetic tree based on *EF1α*. **(D)** A molecular phylogenetic tree based on *Cnnos1*.

**Figure S3.**
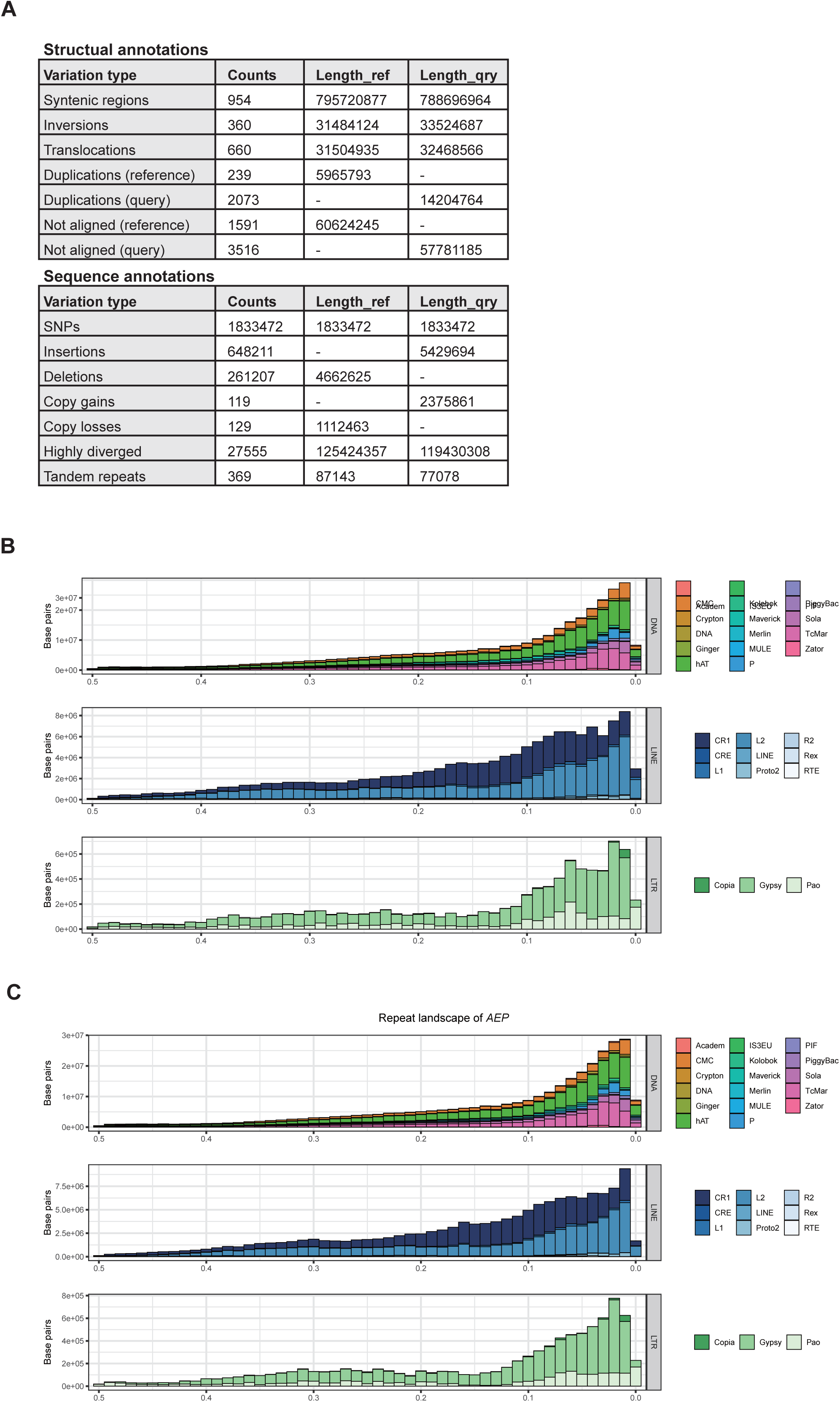
Comparison of the transposable elements between AEP.JNIG and AEP. **(A)** Quantitatively assessed structural and sequence variation identified by SyRI. Structural and sequence variations were identified using SyRI with the AEP assembly as the reference and the AEP.JNIG assembly as the query. Sequence lengths are shown in megabases (Mb). **(B)** Transposable elements (TEs) landscape showing the distribution of repetitive elements in the genome based on Kimura distance. The x-axis represents the Kimura 2-Parameter distance, where lower scores indicate more recent insertions into the genome, while higher scores correspond to older insertions. The y-axis shows the total number of base pairs contributed by each class of transposable elements at each divergence level. The pie chart shows the components and proportions of repetitive sequences in the *H. vulgaris* AEP.JNIG genome. **(C)** Transposable elements (TEs) landscape of *H. vulgaris* AEP, as in B.

**Figure S4.**
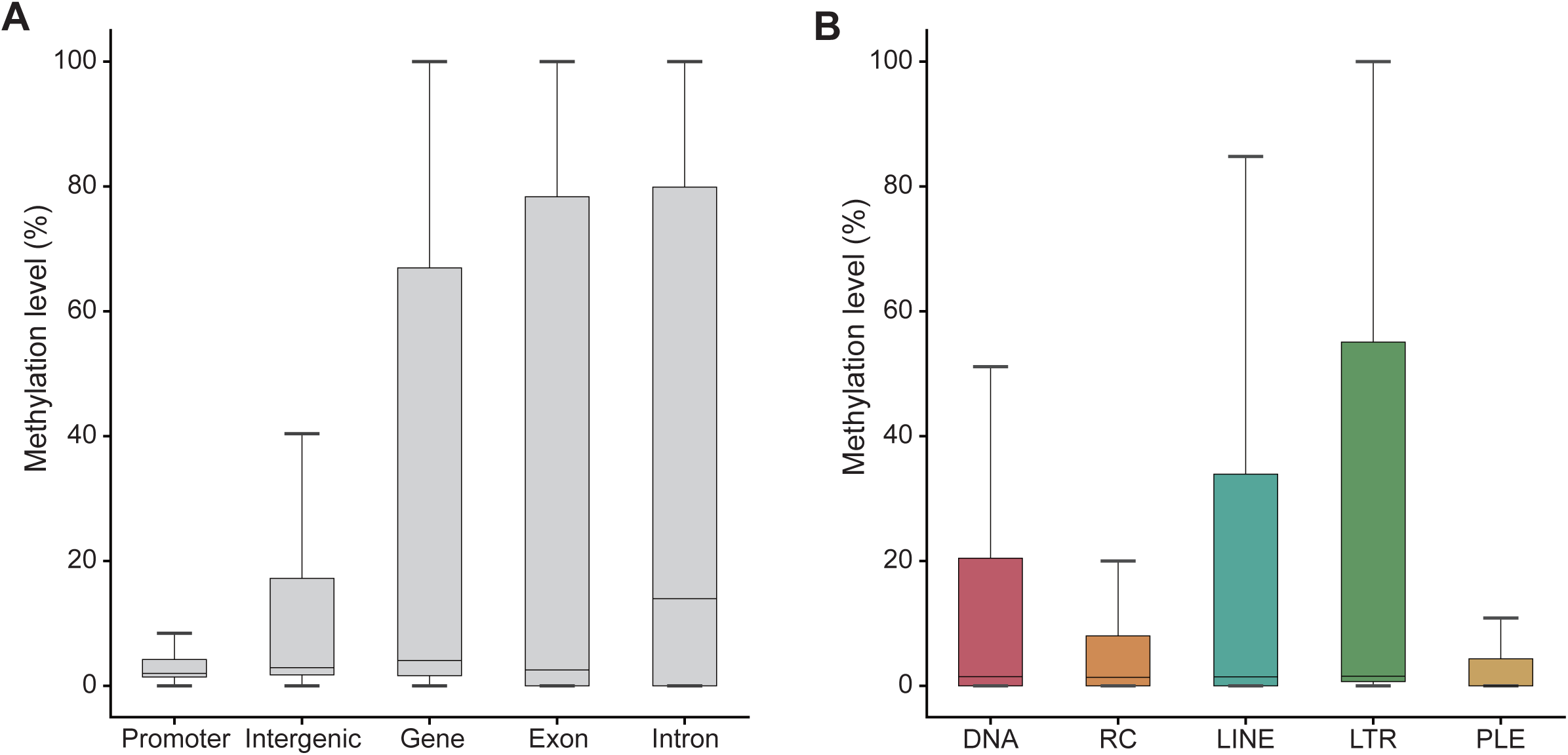
The methylation level of *H. vulgaris* AEP depicted as box plots, related to Fig 3B and 3C. **(A)** The box plot of the methylation level across each genomic region, corresponding to Fig 3B. **(B)** The box plot of the methylation level of each repetitive region, corresponding to Fig 3C.

**Figure S5.**
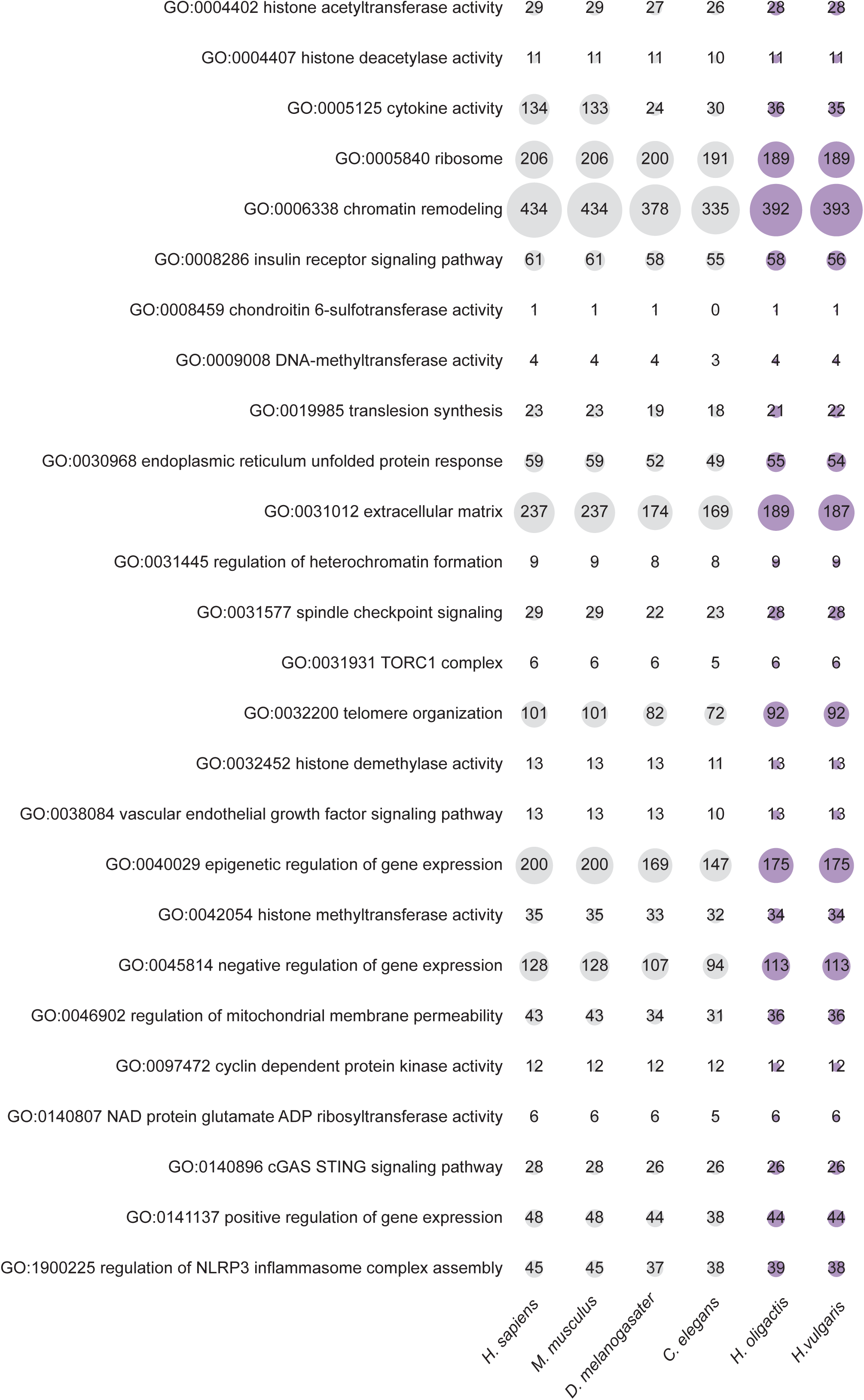
Conservation of aging-related orthologs in *Hydra*, related to Fig 4. Conservation of orthologs related to the *Hallmarks of Aging*, as in Fig 4A. The vertical axis lists representative aging-related pathways (defined as GO terms).

**Figure S6.**
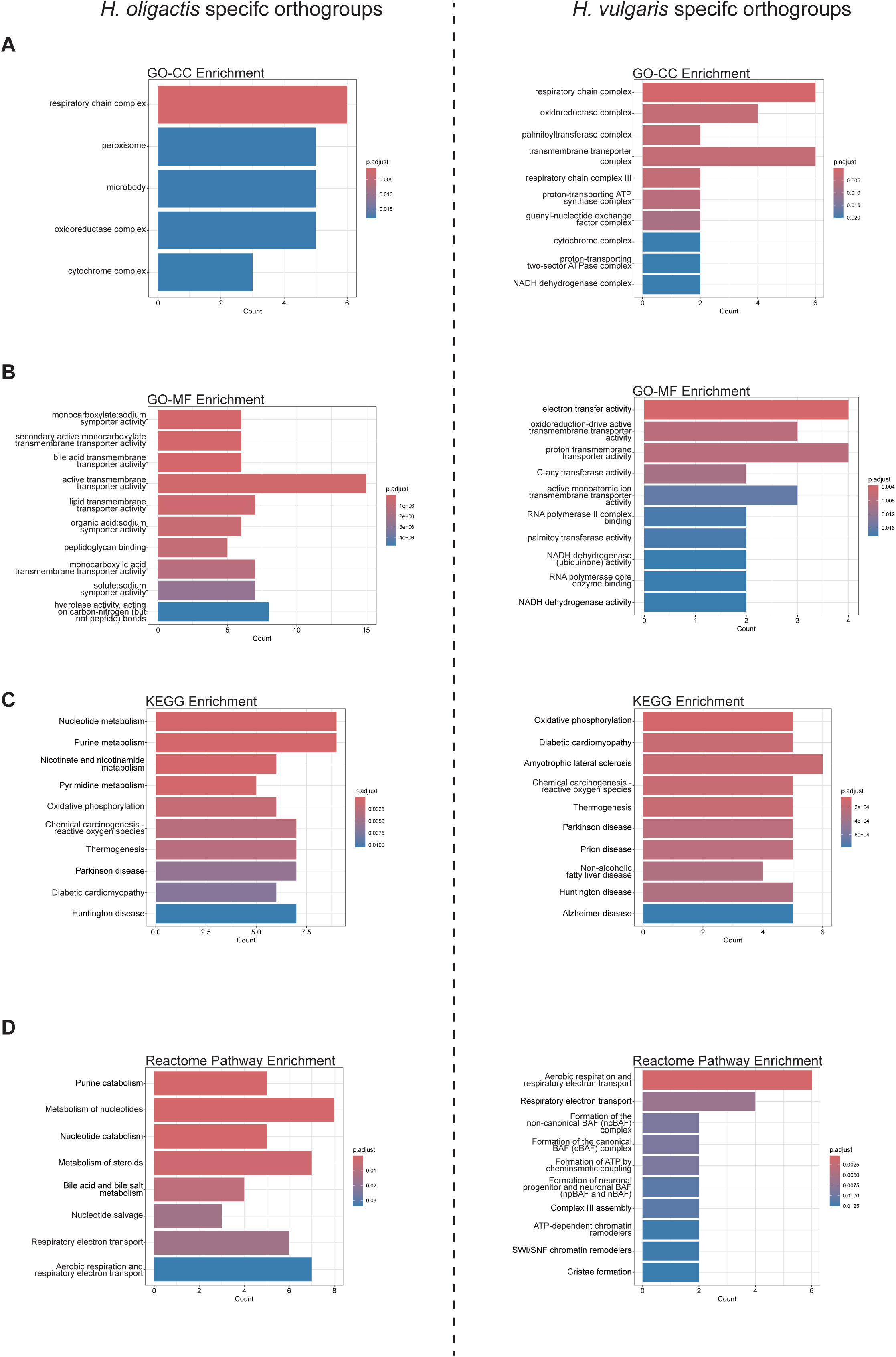
Enrichment analysis results of *H. oligactis*-specific and *H. vulgaris*-specific orthogroups using multiple annotation databases. **(A)** Results of Gene Ontology – Cellular Component (GO-CC) terms. **(B)** Results of Gene Ontology – Molecular Function (GO-MF) terms. **(C)** Results of KEGG pathway terms. **(D)** Results of Reactome pathway terms. In each panel, *H. oligactis*-specific orthogroups are shown on the left, whereas *H. vulgaris*-specific orthogroups are shown on the right.

**Table S1.** Complete list of aging-related genes included in this study. This table provides the full list of aging-related genes identified in this study. For each gene, the second column indicates the associated aging-related pathways, and the third column classifies the gene according to the *Hallmarks of Aging*.

**Table S2.** Summary of the conservation of aging-related genes for each pathway, related to Fig 4 and Fig S5. This table provides an overview of the conservation of aging-related genes identified through GO, KEGG and literature-based searches based on the *Hallmarks of Aging* (López-Otín et al. 2023). The first column lists aging-related pathways, and the subsequent columns show the number of conserved orthogroups in each species.

**Table S3**. Lists of orthogroups specific to *H. oligactis* (65 OGs) and *H. vulgaris* (33 OGs). Each row corresponds to one orthogroup. The second column shows annotations with the corresponding human genes. The third column shows the pathway annotation for each aging-related gene.

**Table S4**. Enrichment analysis of the 33 *H. vulgaris* specific orthogroups and 65 *H. oligactis* specific orthogroups. Significantly enriched terms (adjusted p-value < 0.05) are shown. Ten files are provided, corresponding to all combinations of species-specific gene sets (*H. oligactis*-specific and *H. vulgaris*-specific) and annotation databases (GO biological process, GO cellular component, GO molecular function, KEGG, and Reactome).

